# Divergent cytokine and transcriptional signatures control functional T follicular helper cell heterogeneity

**DOI:** 10.1101/2024.06.12.598622

**Authors:** Lennard Dalit, Chin Wee Tan, Amania A. Sheikh, Ryan Munnings, Carolina Alvarado, Tabinda Hussain, Aidil Zaini, Lucy Cooper, Alana Kirn, Lauren Hailes, Angela Nguyen, Laura Mackay, Nicola Harris, Colby Zaph, Nicole L. La Gruta, Stephen L. Nutt, Kim L. Good-Jacobson, Melissa J Davis, Vanessa L. Bryant, Joanna R. Groom

**Affiliations:** Walter and Eliza Hall Institute of Medical Research; Parkville, VIC, Australia; Department of Medical Biology, University of Melbourne; Parkville, VIC, Australia; Frazer Institute, Faculty of Medicine, The University of Queensland; Brisbane, QLD, Australia; Department of Biochemistry and Molecular Biology, Monash University; Clayton, VIC, Australia; Immunity Program, Biomedicine Discovery Institute, Monash University; Clayton, VIC, Australia; Department of Immunology and Pathology, Monash University; Melbourne, VIC, Australia; Department of Microbiology and Immunology, The University of Melbourne at the Peter Doherty Institute for Infection and Immunity, Melbourne, VIC, Australia; School of Biomedicine, Faculty of Health and Medical Sciences, The University of Adelaide, Adelaide, SA, Australia; Department of Clinical Immunology & Allergy, Royal Melbourne Hospital, Parkville, VIC, Australia

## Abstract

Adaptive immune responses protect against multiple classes of pathogens, including viral, bacterial, fungal, and helminth infections. In all these settings, CD4^+^ T follicular helper (Tfh) cells tailor high-affinity class-switched B cells responses. How Tfh lineage sovereignty is established while allowing for this context-specific functional heterogeneity is unclear. Here, we identify Tfh transcriptional networks in response to diverse infections. While Bcl-6 is the transcriptional linchpin of the core Tfh signature, this is overlayed with pathogen-specific transcriptional modules that shape Tfh function. Cytokine-transcriptional Tfh programing in mouse and human lymphoid tissue demonstrated that type I interferon and TGFβ signaling direct individual Tfh subpopulations to instruct B cell output. Here, we provide a transcriptional map and cell surface resource to interrogate Tfh diversity in humans and mice. This resource can be leveraged to further understand the origins of immune flexibility, perform immune monitoring in infection and antibody-mediated diseases and to develop context-specific vaccines.

**ONE-SENTENCE SUMMARY:** Dalit, Tan and colleagues provide a resource that functionally and transcriptionally profiles T follicular helper cells (Tfh) during diverse pathogen responses to reveal a blueprint for transcriptional flexibility and new tools to interrogate Tfh heterogeneity in mice and humans.

## INTRODUCTION

The adaptive immune system is remarkably flexible. This ensures the clearance of, and generation of memory against, a wide range of pathogen classes, such as viral, bacterial, fungal, and helminth infections^1,2^. Pathogen route, expression of pathogen associated molecular pattern molecules (PAMPs) and antigen affinity and availability are integrated to prompt a tailored response^3–8^. These environmental cues culminate to impact the generation of specialized CD4^+^ T helper (Th) cells that enable pathogen-specific adaptive immunity. Accordingly, Th1 cells produce IFNγ and TNFα to assist during antiviral responses, Th2 cells mediate protection against helminth infection and venoms via production of IL-4, IL-5 and IL-13, and Th17 cells secrete IL-17 family cytokines to protect against bacteria and fungi^1,9,10^. Alongside the differentiation of these Th effector (Teff) cell subsets, CD4^+^ T follicular helper (Tfh) cells are generated to instruct humoral immunity by promoting the germinal center (GC) reaction and differentiation of T-dependent memory B cells and plasma cells, a process dependent on the cardinal Tfh cell cytokine IL-21^11–13^. In contrast to CD4^+^ Teff cells that, despite plasticity, are honed to defend against particular classes of pathogens, Tfh cells drive B cell responses across multiple settings while retaining the ability for context-specific tailoring^1,9,11,14^. Thus, Tfh cells are induced by all categories of pathogens and licensed vaccines, and also influence pathogenesis of diseases such as immunodeficiency, autoimmunity, asthma, allergy, and cancer^11,14^. However, how this exceptional array of Tfh functions is orchestrated is not clear.

Much of the knowledge on Tfh differentiation has focused on the bifurcation model, where Tfh differentiation is contrasted to that of a single Teff helper subset^15^. This work has focused on binary Tfh differentiation in contrast to either Th1, Th2 or Th17 cells^8,16–21^. By emphasizing pathways that are opposed to alternative CD4^+^ differentiation fates, the bifurcation model minimizes the breadth of Tfh functional heterogeneity. Accordingly, additional complexity of this paradigm has been proposed to incorporate Tfh functional diversity^14,40,41^. While multiple cytokine signaling pathways instruct the transcriptional regulators that promote Tfh fate, the role these mediators play in tuning cell function beyond this branch point has not thoroughly been assessed^16,19,22–39^ and thus a unifying understanding of how Tfh responses are tailored in response to pathogen-specific cues is lacking.

Evidence of Tfh heterogeneity has been detailed by distinct expression of transcriptional regulators, varied production of Th cytokines and chemokine receptor surface expression^14,40^. CD4^+^ T cell lineage defining-transcription factors act to upregulate cell identity factors, and to repress alternative fates. Despite this, it is increasingly appreciated that the Tfh lineage-defining transcription factor, B-cell lymphoma-6 (Bcl-6) can be co-expressed with transcription factors that define Th1, Th2 and Th17 differentiation, namely T-bet, GATA3, and RORγt^11,14,29,40–43^. Aligned with borrowed transcription factors, Tfh cells can produce IFN-γ, IL-4, IL-10 and IL- 17 cytokines indicative of other Teff cell subsets^32,44,45^. In addition to production of IL- 21, Teff cytokines tailor B cell responses and dictate the class-switch isoform of high-affinity antibodies^12,13,19,40,44,46^. In humans, subpopulations of circulating blood memory Tfh (cTfh) cells with specialized B helper properties are identified via the expression of chemokine receptors, where CXCR3^+^ cTfh cells resemble Th1, CXCR3^-^CCR6^-^ cells resemble Th2, and CCR6^+^ cells resemble Th17 cells, respectively^14,47–49^. Immune profiling of specific cTfh populations correlates with disease and protective vaccine outcomes^11,49–53^. However, the use of inflammatory chemokine receptors to define cTfh cells may reflect cell environment rather than identity, and there is limited evidence of CXCR3^+^, CCR6^+^ and CXCR3^-^CCR6- populations within human tonsil^54^. Thus, although Tfh cells can parallel Teff on multiple levels, the extent of Tfh heterogeneity or how distinct Tfh subpopulations are established remains unclear^40^.

To address these questions, we profiled the spectrum of functional Tfh cell states induced by diverse pathogens. We establish a core Tfh signature, that is centrally regulated by Bcl-6, independent of Th effector transcriptional regulators and contains previously unknown regulators of Tfh biology. We reveal that, beyond Tfh identity, diverse cytokine-guided transcriptional modules enable tailoring of Tfh cell functional capacity. These cytokine signaling pathways were evident in human tonsil, where individual Tfh populations can be identified by distinct cell surface molecules.

Combined we present a transcriptional and cell surface resource to understand the mechanics of Tfh heterogeneity and plasticity, reveal opportunities to fine-tune humoral responses for vaccination and to understand Tfh dynamics during infection, vaccination, antibody deficiency and antibody-mediated disease.

## RESULTS

### Functionally heterogenous Tfh populations are induced by diverse pathogen infections

The developmental pathway towards Tfh generation relies on the integration of TLR signaling and dendritic cell activation which are dictated by pathogen class, route of infection and draining lymph node^2,3,10,55,56^. Considering this, we hypothesized that pathogen-specific cues result in functionally distinct Tfh subpopulations, each having distinct capacity to shape B cell responses. To test this, we investigated Tfh and Teff differentiation and function generated during viral, helminth, and bacterial infections. As Tfh cells evolve over time^46,57,58^, the early peak of Tfh cell accumulation, defined by Tfh and GC B cell frequency, was individually assessed for each infection, assessing polyclonal Tfh (CD3^+^CD4^+^CD44^+^CXCR5^+^Ly6C^-^) and Teff (CD3^+^CD4^+^CD44^+^CXCR5^-^Ly6C^+^) cell compartments in viral (lymphocytic choriomeningitis virus (LCMV, acute Armstrong strain), Influenza A (Influenza, HKx31 H3N2 strain), helminth (*Trichuris muris* (*T. muris*), *Heligmosomoides polygyrus* (*H. polygyrus*)), and bacterial (*Citrobacter rodentium* (*C. rodentium*)) infections (Figures 1A, S1A, and S1B). Rather than minimizing their influence, we analyzed the natural infection route and draining lymph node for each infection (inguinal, brachial and axillary lymph nodes for LCMV; mediastinal and cervical lymph nodes for Influenza A; and mesenteric lymph node for remaining pathogens) to investigate the breadth of Tfh heterogeneity^59^. Tfh cells were generated to a varying degree in all infections, as reflected by the frequency of Tfh cells within the CD4^+^CD44^+^ cell population and the ratio between Tfh and Teff populations (Figures 1B and 1C).

**Figure 1.**
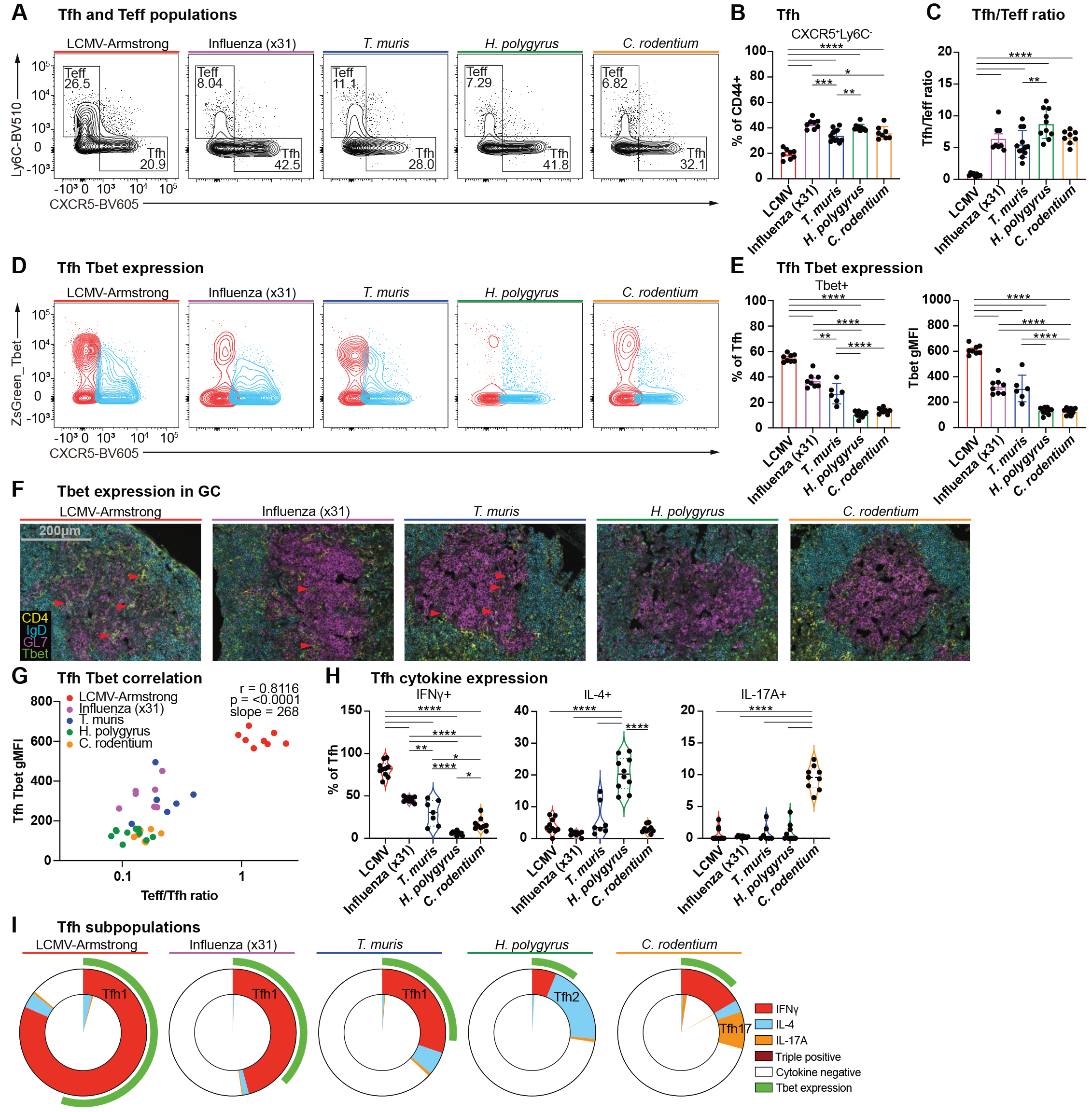
Functionally heterogenous Tfh populations are induced by diverse pathogen infection. Analysis of (A to I) draining lymph node cells from (A to C, H and I) wildtype and (D to G) ZsGreen_T-bet reporter mice infected with indicated pathogens at early peak GC response. (A) Representative plots (B) frequency of CD4^+^CD44^+^CXCR5^+^Ly6C^-^ Tfh cells and CD4^+^CD44^+^CXCR5^-^Ly6C^+^ Teff cells, and (C) ratio of Tfh/Teff cells. (D) Representative plots, (E) ZsGreen_T-bet reporter^+^ frequency and gMFI of CD4^+^CD44^+^CXCR5^+^Ly6C^-^ Tfh cells. (F) Immunofluorescent staining of draining lymph node GCs. Red arrows indicate ZsGreen_T-bet reporter^+^ Tfh cells. Yellow: CD4, blue: IgD, magenta: GL7, green: ZsGreen_T-bet reporter. Scale bar 200 μm. (G) Correlation of ZsGreen_T-bet reporter expression to ratio of Teff/Tfh cells across infections. (H) Frequency of Tfh cells produced IFNγ, IL-4, IL-17A. (I) IFNγ^+^ Tfh1, IL- 4^+^ Tfh2, and IL-17^+^ Tfh17 cells displayed as parts of whole Tfh population. Inner slice displays cytokine co-expression. Outer ring (green) indicates proportion of ZsGreen_T-bet^+^ Tfh expression.

The Th1 lineage defining-transcription factor, T-bet (encoded by *Tbx21*) plays a context-specific role in Tfh bifurcation and function in response to viral infections^19,42,60^. Accordingly, we sought to investigate the expression of the ZsGreen-T-bet reporter across diverse pathogen classes^61^. Teff cells reported higher T-bet than Tfh cells in all infections (Figures 1D and S1C). T-bet expression level was graded within the Tfh cell compartment across infections (Figure 1E). While viral infections induced the highest Tfh T-bet expression, the helminth *T. muris* expressed comparable levels to that seen in Influenza, with Tfh cells generated during *H. polygyrus* and bacterial *C. rodentium* infections reporting low T-bet expression (Figure 1E). Confocal analysis of draining lymph nodes displayed conserved GC (GL7^+^IgD^-^) morphology, in line with the relative frequency of Tfh and GC B cells across all pathogen classes^11,14^ (Figure S2). Within GC structures, we identified T- bet^+^ Tfh cells in LCMV, Influenza, and *T. muris* infected tissue, but not from *H. polygyrus* and *C. rodentium*, consistent with the higher level of Tfh T-bet expression in these infections (Figure 1F). As LCMV and *T. muris* reported the highest Tfh T-bet expression and were the least skewed towards Tfh generation (Figure 1B), this confirmed the relationship between T-bet and Tfh/Teff bifurcation across multiple infection types^19^. Indeed, higher T-bet expression correlated with a Teff-skewed T cell response (r=0.8116) demonstrating the context-specific role for T-bet extends beyond viral pathogens to helminth and bacterial infections (Figure 1G)^8,24^.

As Tfh cells direct B cell responses via the production of cytokines and the engagement of cell surface receptors ^11,40,44,62^, we next investigated how diverse pathogens influence the cytokine potential of Tfh cells. Each pathogen-induced Tfh population displayed a distinct cytokine profile (Figures 1H and S3A). Mirroring Teff cytokine production for each infection, Tfh cells induced by viral LCMV and Influenza infections expressed the highest level of IFNγ, along with *T. muris* helminth infection, whereas Tfh cells induced by *H. polygyrus* and *C. rodentium* expressed the highest levels of IL-4 and IL-17A respectively (Figures 1H and S3B). The majority of Tfh cells demonstrated cytokine specialization, with minimal cells producing IFNγ, IL-4, or IL- 17 in combination (Figure 1I). Intracellular cytokine expression was confirmed using IFNγ-GFPx4C13R reporter mice for *T. muris* infection which, in addition to similar IFNγ and IL-4 expression, indicated the presence of an IL-13-expressing Tfh population (Figure S3C)^63^. Serum cytokine analysis largely reflected that observed in draining lymph nodes, with additional expression of TNFα seen in each infection, suggesting that Tfh cytokine production reflects the overall infection milieu (Figure S3D). Further, overlaying Tfh T-bet expression onto Tfh cytokine production revealed that beyond Tfh/Th1 bifurcation, T-bet underlies the IFNγ^+^ Tfh1 subpopulation across all infections (Figure 1I)^19,60^.

### Pathogen-induced Tfh heterogeneity correlates with tailored B cell responses

We next investigated how the variation in Tfh cytokine expression correlated with GC B cell and antibody production. GC B (B220^+^CD138^-^IgD^lo^CD38^-^CD95^+^) cells were quantified, with *H. polygyrus* and *T. muris* generating the largest GC B cell response (Figures 2A and S4A). Although *H. polygyrus* showed the highest Tfh cell-bias, the frequency of GC B cells and Teff/Tfh ratio were not inversely correlated across all infections (r=-0.3483) (Figure S4B). Memory B (MBC, B220^+^CD138^-^ IgD^lo^CD38^+^CD95^lo^) cell analysis revealed that LCMV induced the most memory B cells, followed by Influenza and *C. rodentium* (Figure 2B). The relative frequency of MBCs correlated with a high Teff/Tfh ratio (r=0.7149) as well as high Tfh T-bet^+^ expression (r=0.8197), that was largely driven by LCMV (Figure 2B). Further, serum antibody analysis demonstrated distinct IgG class switch isotype expression, consistent with each infection producing a unique combination of Tfh cytokines (Figure 2C). Infections that expressed the highest Tfh T-bet^+^ expression, LCMV, *T. muris*, and Influenza induced pro-inflammatory IgG2b. In Influenza infection this was also accompanied by IgG3 and IgG1. In contrast, *H. polygyrus* produced the highest IgG1, while *C. rodentium* led to high IgG2b production, with intermediate production of IgG1 and IgG3. Combined, diverse pathogen infections induce distinct populations of Tfh cells that exhibit distinct cytokine profiles which correlate to GC and MBC output and distinct antibody isotype usage. Further, we observe that Tfh cell and GC heterogeneity does not entirely align within pathogen class, such that Tfh cells arising during *T.muris* infection were more similar to those in LCMV or Influenza than *H.polygyrus* infection.

**Figure 2.**
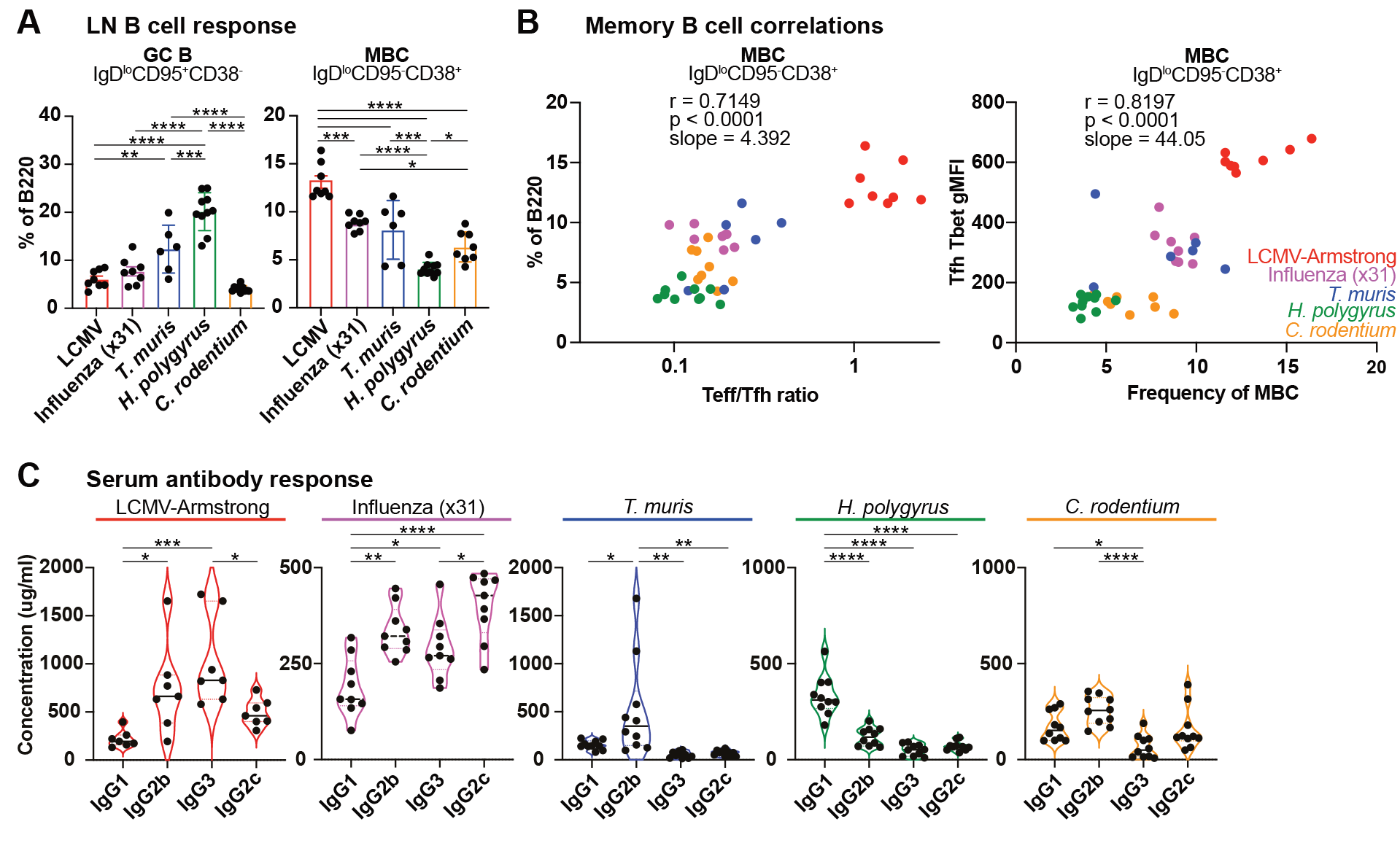
Diverse pathogen infection demonstrate tailored B cell responses. Analysis of (A to C) draining lymph node cells and (D) serum from (A and C) wildtype and (B) ZsGreen_T-bet reporter mice infected with indicated pathogens at early peak GC response. (A) Analysis of B220^+^IgD^lo^CD95^+^CD38^-^ GC B cells and B220^+^IgD^lo^CD95^lo^CD38^+^ MBCs. (B) Correlation of frequency of MBC to the ratio of Teff/Tfh cells and to ZsGreen_T-bet reporter expression gMFI across infections. (C) Serum IgG isotype concentration. Data show experiments of 6-10 mice per group and mean ± SEM. Statistical tests: one-way ANOVA of multiple comparisons and Pearson correlation with two-tailed P value.

### The conserved Tfh signature is independent of Th lineage defining transcription factors

The bifurcation paradigm for Tfh differentiation, compared to individual alternate Teff fates, has been central to our understanding of Tfh transcriptional programs^15^. As diverse pathogens induce functionally distinct Tfh subpopulations that correlated with GC output, this binary comparison of CD4^+^ cell differentiation is insufficient to identify a core Tfh program that is centrally induced regardless of pathogen class. To address this, we performed RNAseq on GC T cell populations, analyzing Tfh (CD4^+^CD44^+^CXCR5^hi^PD1^hi^) cells compared to Teff (CD4^+^CD44^+^CXCR5^-^PD1^-^) cells at the early peak of GC formation in LCMV, Influenza, *T. muris*, *H. polygyrus*, and *C. rodentium* (Figure S5A). As IL-21^+^ cells have been shown to exhibit increased GC residency (referred to as GC-Tfh) with distinct transcriptional profiles, we used the IL- 21^GFP^ transcriptional reporter to profile both IL-21^+^ and IL-21^-^ Tfh cells^12,64^. Further, we isolated T follicular regulatory (Tfr, FoxP3^+^CXCR5^+^PD1^+^) cells, a subset that represses GC size and facilitates the emergence of high-affinity GC B cells^58,65,66^.

Principal component analysis (PCA) of all samples, separated firstly by cell type on PC1 (32.34%) and PC2 (21.18%), with IL-21^+^ Tfh and IL-21^-^ Tfh cells being more related to either Tfr and Teff samples (Figure S5B). Secondary to cell type, samples separated based on infection type on PC3 (7.03%) and PC4 (4.44%) (Figure S5B). In contrast to previous studies, differential expression gene analysis of IL-21^+^ Tfh and IL-21^-^ Tfh cells revealed they were transcriptionally similar, with only 3 key genes *Ly6c2, Il7r* (encoding CD127), and *Sell* (encoding CD62L) downregulated in the IL-21^+^ Tfh population in 4 of 5 infections (Figure S6A). Cell surface staining confirmed the regulation of these factors at the protein level in LCMV and Influenza (Figures S6B to S6G). Given the strong transcriptional identity, both IL-21^+^ Tfh and IL-21^-^ Tfh populations were used to define the core Tfh transcription program. For this, we combined each of the Tfh samples from the 5 infections and performed differential expression analysis relative to Teff from each infection (Figure 3A).

**Figure 3.**
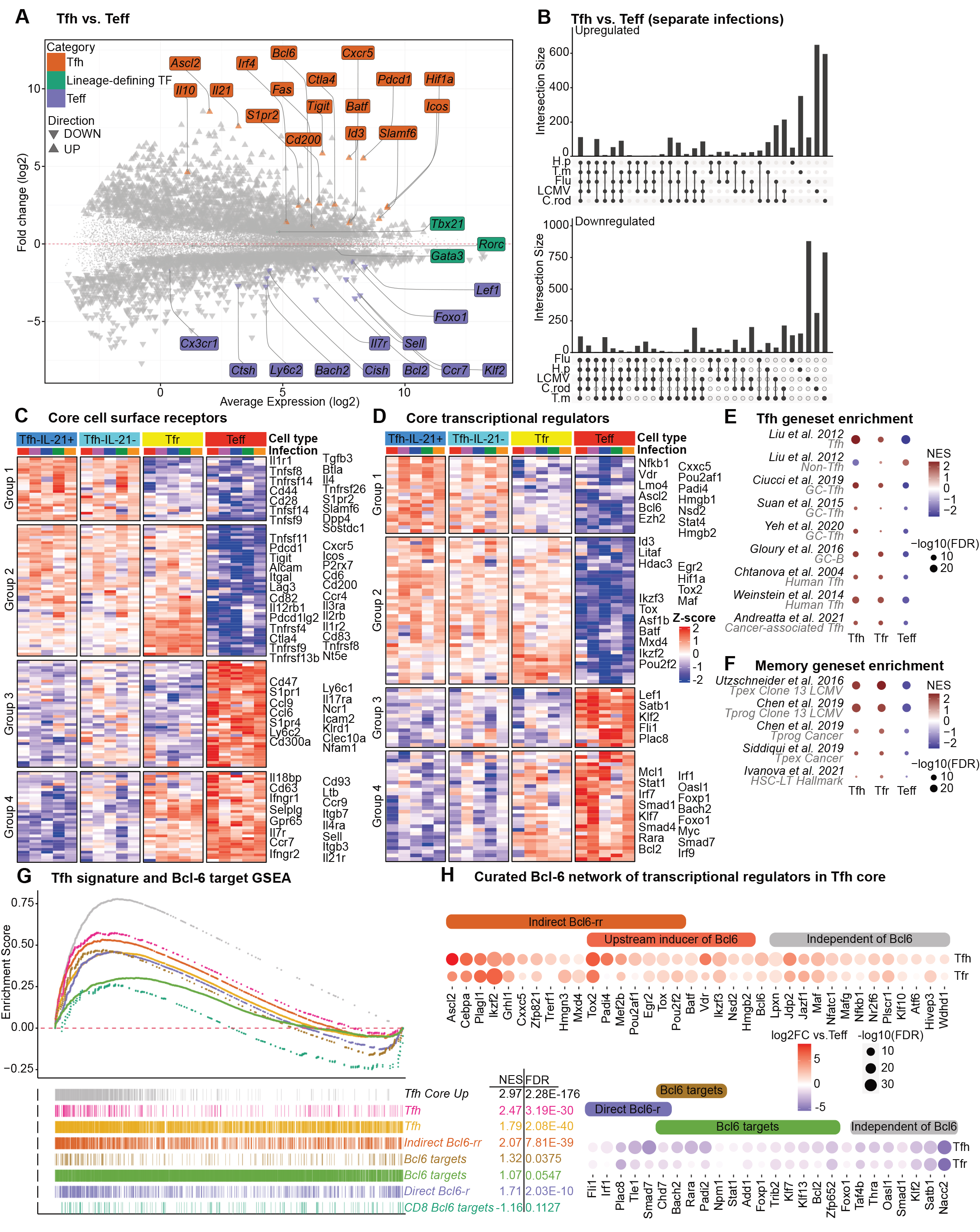
Conserved Tfh signature is independent of lineage defining transcription factors and revolves around Bcl-6 as the core regulatory linchpin. (A to H) RNAseq of sorted CD4^+^CD44^+^PD1^+^CXCR5^+^FoxP3-RFP^-^IL-21-GFP^+^ Tfh, CD4^+^CD44^+^PD1^+^CXCR5^+^FoxP3-RFP^-^IL-21-GFP^-^ Tfh, CD4^+^CD44^+^PD1^+^CXCR5^+^FoxP3-RFP^+^IL-21-GFP^+^ Tfr, and CD4^+^CD44^+^PD1^-^CXCR5^-^FoxP3-RFP^-^IL-21-GFP^-^ Teff cells from draining lymph nodes of mice infected with the pathogens described in Fig 1. (A) MA-plot visualizing log fold-change against the mean expression of genes expressed differentially between Tfh and Teff populations for five infections. (B) Upset plot showing intersections of genes expressed differentially between Tfh and Teff populations for each infection. Common DE genes across 3 or more infections used to derive the core Tfh signature. False discovery rate (FDR) < 0.05. (C and D) Heatmap of core Tfh signature genes (row-based z-score of normalized log2 counts per million) for (C) cytokine and cell surface receptor genes and (D) transcriptional regulator genes. (E and F) GSEA of differentially expressed genes in Tfh (Tfh vs. Teff comparison), Tfr (Tfr vs. Teff), and Teff cells (Teff vs. Tfh) for all infections. (E) Normalized enrichment score plots (NES) of enrichment for Tfh, non-Tfh, GC-Tfh, GC-B, human Tfh, and cancer-associated Tfh cell programs and (F) precursor of exhausted (Tpex), exhausted progenitor (Tprog), and long-term hematopoietic stem cell (HSC-LT) gene programs in Tfh, Tfr, and Teff populations. NES score represents enrichment of genes (sets) relative to each comparison, correcting for multiple testing. (G and H) GSEA of differentially expressed genes in Tfh (Tfh vs. Teff contrast) and Tfr cells (Tfr vs. Teff contrast) combined for five infections. (G) Barcode and ES plots displaying enrichment of genes (ES and NES) upregulated in core Tfh signature (Tfh Core Up), Tfh genes, genes indirectly regulated by Bcl-6 through repressor-of-repressor circuits (Indirect Bcl6-rr), genes directly repressed by Bcl-6 (Direct Bcl6-r), and Bcl-6 target genes (Bcl6 targets) in Tfh cells (Tfh vs. Teff) contrast^87–90,101^. (H) Bubble plot of Bcl- 6 network transcriptional regulator genes in Tfh core displaying log2FC differences of Tfh and Tfr populations. The size of the bubble represents -log10(FDR) and the color indicates log2FC compared to Teff cells. Colored bar indicates genes present in Tfh core and Bcl-6 gene sets from Figure 2G. Novel genes proposed to be independent of Bcl-6 network indicated with grey bar. Data show independent samples of 2-3 per cell type per infection.

Validating this approach, known Tfh genes (*Bcl6*, *Cxcr5*, *Pdcd1* (encoding PD1), and *Cd200*) were upregulated and known Teff genes (*Klf2*, *Ccr7*, *Il7r*, *Ly6c1* and *Sell*) were downregulated (Figure 3A). While Th lineage defining factors *Tbx21*, *Gata3*, and *Rorc* have each been shown to mediate the bifurcation between Tfh cells and their respective Th subsets, the core Tfh signature was independent of these factors, eliminating them as central determinants of Tfh differentiation (Figure 3A). In contrast, the transcriptional regulators *Foxo1*, *Bach2 and Foxp1* were present in the Teff core signature, indicating their potential to act in opposition to the Tfh program regardless of the pathogen encountered^67–73^. We next performed pairwise differential expression analysis of Tfh versus Teff for each infection to identify genes common across 5, 4 or 3 infections (Figure 3B). From this we defined a core Tfh signature that contained genes from Tfh versus Teff analyses from each infection, that showed common significant directional regulation across 3 or more infections (719 upregulated and 810 downregulated genes).

Analysis of transcript expression of genes encoding cell surface proteins, cytokine signaling components and transcription factors revealed the core Tfh signature fell into 4 groups; 1. Tfh core genes down-regulated by both Teff and Tfr; 2. Tfh genes co-expressed in Tfr cells, 3. Teff core genes down-regulated by Tfh and Tfr; and 4. Teff genes co-expressed in Tfr cells (Figures 3C and 3D). Each of the Tfh core genes (comprising Groups 1 and 2) contain known and novel regulators of Tfh identity, fate and function. Genes with the largest normalized expression differences in Group 1 highlighted Tfh function and receptors that promote GC positioning and B cell interactions (*S1pr2*, *Tnfsf8* (encoding OX40), *Btla*, *Il4*)^74–76^, genes that assist in the transition from Tfh to Tfr (*Sostdc1*)^77^, transcriptional and epigenetic regulators of cell fate (*Bcl6*, *Ascl2*, *Ezh2*)^11^, along with genes not previously associated with Tfh identity or function (*Dpp4*, *Padi4*, *Lmo4*). Group 2 genes established a shared Tfh and Tfr signature, which contained genes required for Tfh and Tfr identity (*Cxcr5*, *Pdcd1*, *Id3*, *Icos*, *Cd200*, *Hif1a*), along with genes suggestive of memory potential (*Tox*, *Tox2, Hdac3, Mxd4)*^78–81^. In contrast, the Teff core signature (comprising Groups 3 and 4) contained antagonists to Tfh lineage commitment (*Foxo1, Bach2*, *Foxp1*)^67–73^, T cell zone positioning receptors (*Ccr7*, *Sell*, *S1pr1*), inflammatory mediators and signaling (*Ly6c1*, *Ltb, Ifngr2*, *Irf7*, *Oasl1*, *Smad7*), and proliferation and apoptosis inhibitors (*Mcl1*, *Bcl2*, *Myc*). Further, gene-set enrichment analyses (GSEA) with published datasets with the normalized enrichment score (NES) confirmed that the Tfh core aligns with GC-Tfh cells^64,82,83^, GC-B cells^84^ and additional infection-, vaccine- and cancer-induced Tfh cells in mice and humans (Figure 3E). In line with the observation that both Tfh and Tfr cells (Group 2 genes) expressed checkpoint receptors (*Pdcd1*, *Ctla4*, *Lag3*, *Tigit*) and the aforementioned drivers of long-term memory, GSEA-NES analysis revealed alignment with gene signatures of CD8^+^ T cell precursor and progenitors of exhausted cells (Tpex and Tprog) from chronic infection and cancer and signatures associated with long-term hematopoietic stem cell memory, relative to Teff cells (Figure 3F). Thus, the core Tfh signature is suggestive of increased memory potential, similar to that observed for stem-like CD8^+^ memory cells^85,86^.

### Bcl-6 commands the core Tfh signature

Bcl-6 acts to promote multiple aspects of Tfh fate and function^11,16–18^. Therefore, we used GSEA to investigate Bcl-6 target binding relative to the Tfh core signature. As expected, published Tfh cell expression datasets were enriched to the Tfh core signature (Figure 3G)^87,88^. Additionally, the core signature was enriched for Bcl-6 occupancy loci from mouse and human Tfh cell ChIPseq data, although these data were not selected for activation or repressive marks^89,90^. As Bcl-6 primarily acts as a repressor of gene expression^11^, the core Tfh signature was more significantly aligned to indirect targets (repressor-of-repressor, Bcl6-rr) than direct targets (regulated, Bcl6-r), and direct Bcl-6 targets in CD8^+^ T cells (Figure 3G)^87^. We next investigated how individual transcription factors expressed by both Tfh and Teff cells were influenced by Bcl-6. In line with the role of Bcl-6 as an obligate repressor that represses non-Tfh fate, genes that were direct Bcl-6 targets had decreased expression in Tfh and Tfr cells (Figure 3H)^16,17^. Genes that are indirectly regulated by Bcl-6 through repressor-of-repressor circuits had increased expression in Tfh and Tfr cells. Additionally, Bcl6-rr genes overlapped with genes known to be upstream inducers of Bcl-6. While a small group of transcriptional regulators were separate from known Bcl-6 circuits, the majority were within the Bcl-6 network, emphasizing that Bcl-6 underpins Tfh differentiation, in a manner that is independent of pathogen class.

### Graded expression of lineage defining transcription factors overlay the core Tfh signature to tailor Tfh heterogeneity

While T-bet expression correlated with IFNγ^+^ Tfh1 cells (Figure 1I), other regulators of Tfh heterogeneity remain unknown. To investigate this, we examined the pathogen-specific transcriptional programs of Tfh cells across infections.

Multidimensional Scaling (MDS) analysis indicated pathogen-directed Tfh cells separated by infection (Figures 4A and S5B). To establish pathogen-specific signatures, we conducted differential expression analysis and compared samples from each infection to the combined transcriptomes of all others, for both Tfh and Teff populations. In Influenza and *H. polygyrus* a large proportion of pathogen-specific genes were shared between Tfh and Teff cells, while this was not the case for LCMV or *T. muris* (Figure 4B). Given this variability, we defined pathogen-specific Tfh signatures to include the shared and Tfh distinct differentially regulated genes.

**Figure 4.**
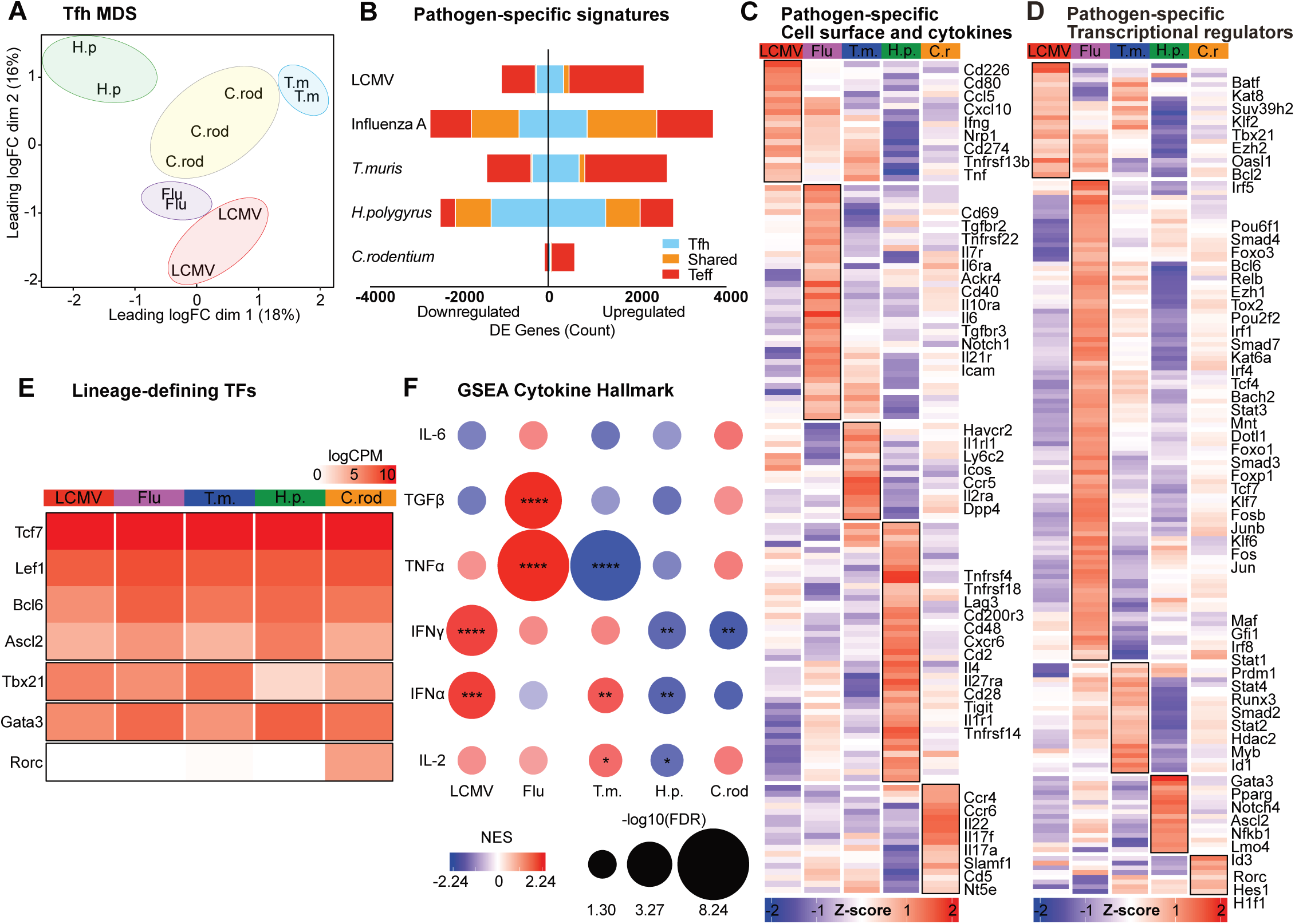
Identification of Pathogen-specific Tfh heterogeneity. RNAseq of sorted CD4^+^CD44^+^PD1^+^CXCR5^+^FoxP3-RFP^-^IL-21-GFP^+^ Tfh, CD4^+^CD44^+^PD1^+^CXCR5^+^FoxP3-RFP^-^IL-21-GFP^-^ Tfh cells from draining lymph nodes of mice infected with the indicated pathogens (as in Fig 1). (A) Multi- dimensional scaling (MDS) plot of CD4^+^CD44^+^PD1^+^CXCR5^+^FoxP3-RFP^-^IL-21-GFP^+^ Tfh transcriptomes showing the separation of samples by infection type along dim 1 and 2. (B) Genes expressed differentially for Tfh (single infection vs. all other infections) and Teff cells (single infection vs. all other infections) (FDR < 0.05). Orange: common pathogen-specific DE genes in both Tfh and Teff cells, blue: pathogen-specific signatures unique to Tfh cells, red: pathogen-specific signatures unique to Teff cells. (C to D) Heatmaps of pathogen-specific Tfh signature genes (average row-based z-score of normalized log2 counts per million) for selected (C) cytokine and cell surface receptor genes and (D) transcriptional regulator genes. (E) Heatmap of lineage-defining transcription factors (log2 counts per million) for each indicated infection. (F) GSEA of pathogen-specific signatures using MSigDB “Hallmark” gene sets for cytokine signaling and response. The size of the bubble represents -log10(FDR) and the color indicates NES. Data show independent samples of 2-3 per cell type per infection.

Analysis of cell surface and cytokine genes revealed Tfh from each pathogen provided specialized help via distinct cytokine and chemokine (*Ifng*, *Cxcl10*, *Il6*, *Il4*, *Il22*, *Il17*), and inhibitory and co-stimulatory molecules (*Cd226* (encoding DNAM), *Cd274* (encoding PD-L1), *Cd80*, *Tnfrsf13b* (encoding TACI), *Tnfrsf22* (encoding TRAIL), *Cd40*, *Havcr2* (encoding TIM3), *Icos*, *Tnfrsf4* (encoding OX40), *Cd28*, *Tigit*) (Figure 4C). Tfh cells arising from each infection also expressed a distinct set of transcriptional and epigenetic regulators, suggesting directed differentiation of Tfh subpopulations beyond Tfh/Teff bifurcation (Figure 4D). Interestingly, due to differential expression inclusion criteria, some genes identified in the core Tfh signature were additionally found within individual pathogen-associated signatures (including *Bcl6*, *Ascl2*, *Ezh2*,*Tox2*, *Stat4*), indicating additional tailoring of gene expression within individual Tfh subpopulations. We observed the lineage defining transcription factors *Tbx21*, *Gata3* and *Rorc* were differentially expressed within the pathogen-specific signatures, in contrast to Bcl-6 and others in the Tfh core (Figures 4D and 4E). Visualizing gene expression as normalized log counts per million reads (logCPM) indicated that graded expression of *Tbx21* and *Gata3* were reciprocally expressed between infections, while *C. rodentium* was the only infection that induced Tfh cell *Rorc* expression (Figure 4E).

### Pathogen-specific Tfh heterogeneity is instructed by distinct cytokine signaling

We next performed enrichment analysis based on the MSigDB database’s cytokine Hallmark gene sets to understand the involvement of cytokine signaling pathways that may induce distinct Tfh subpopulations. Type I and type II interferon (IFN-I and IFN-II) signaling genes were positively enriched in Tfh cells from LCMV and *T. muris* (Figure 4F). In contrast, these pathway genes were negatively enriched for *H. polygyrus* and *C. rodentium*, indicating a key role for IFNs as drivers of pathogen-specific Tfh flexibility. Despite being a viral infection, the decreased IFN-I signaling observed in Tfh cells from Influenza was consistent with the generation of Tfh-skewed responses, compared to those in LCMV (Figures 1C and 4F)^26^. Instead, Influenza was enriched for TGFβ signaling. Although *in vitro* Tfh differentiation is assisted by TGFβ, it is non-essential or inhibitory for mouse Tfh cells^16,33–36^.

However, in murine Influenza infection, TGFβ signaling promotes Tfh formation^37^. We reasoned this context-specific role for TGFβ signaling was consistent with the concept of Tfh diversity, whereby distinct pathogen-specific cues induce specific Tfh subpopulations. Similarly, while IFN-I signaling promotes Th1 differentiation at the expense of Tfh cells, depending on the context, IFN-I can also promote Tfh^26–28^.

Thus, we next investigated the role of TGFβ and IFN-I signaling in the infection models that were dominated by these cytokine pathways, Influenza and LCMV respectively. For this, Tfh subpopulations were categorized into Tfh1, Tfh2, and Tfh17 subpopulations according to CXCR3 and CCR6 chemokine profiles, as IFNAR deficiency had an extraneous effect on IFNγ expression. Consistent with previous work, targeted T cell deletion of TGFβR2 (*Tgfbr2*^fl/fl^Cre^LCK^) decreased the Tfh/Teff ratio (Figure S7A). Beyond this, the quality of Tfh cells produced was altered, with increased Tfh1 and decreased Tfh2 and Tfh17 populations (Figure 5A). Altered Tfh subpopulations impacted the B cell response, with decreased IgG1^+^ and reciprocal increase evident on IgG2c^+^ GC B cells (Figure 5B and 5C). Additionally, GC cycling was dysregulated with an increased dark zone (CD86^-^CXCR4^+^) and decreased light zone (CD86^+^CXCR4^-^) GC populations (Figures 5B and S7B). While IFNAR deficiency (*Ifnar*^-/-^) resulted in the reciprocal alteration in Tfh/Teff generation to TGFβR2 deficiency (Figure S7A), within Tfh subpopulations, loss of IFN-I signaling had an opposite effect, with decreased Tfh1 and increased Tfh2, also with decreased Tfh17 subpopulations (Figure 5D). Altered Tfh subpopulations were reflected in the B cell response with increased IgG1^+^ GC B, decreased dark zone GC B cells, and increased serum IgG1 concentration (Figures 5E and 5F). Thus, beyond the influence of Tfh and Teff bifurcation, distinct pathogen-induced cytokine environments determine the capacity of Tfh cells to direct B cell and GC output.

**Figure 5.**
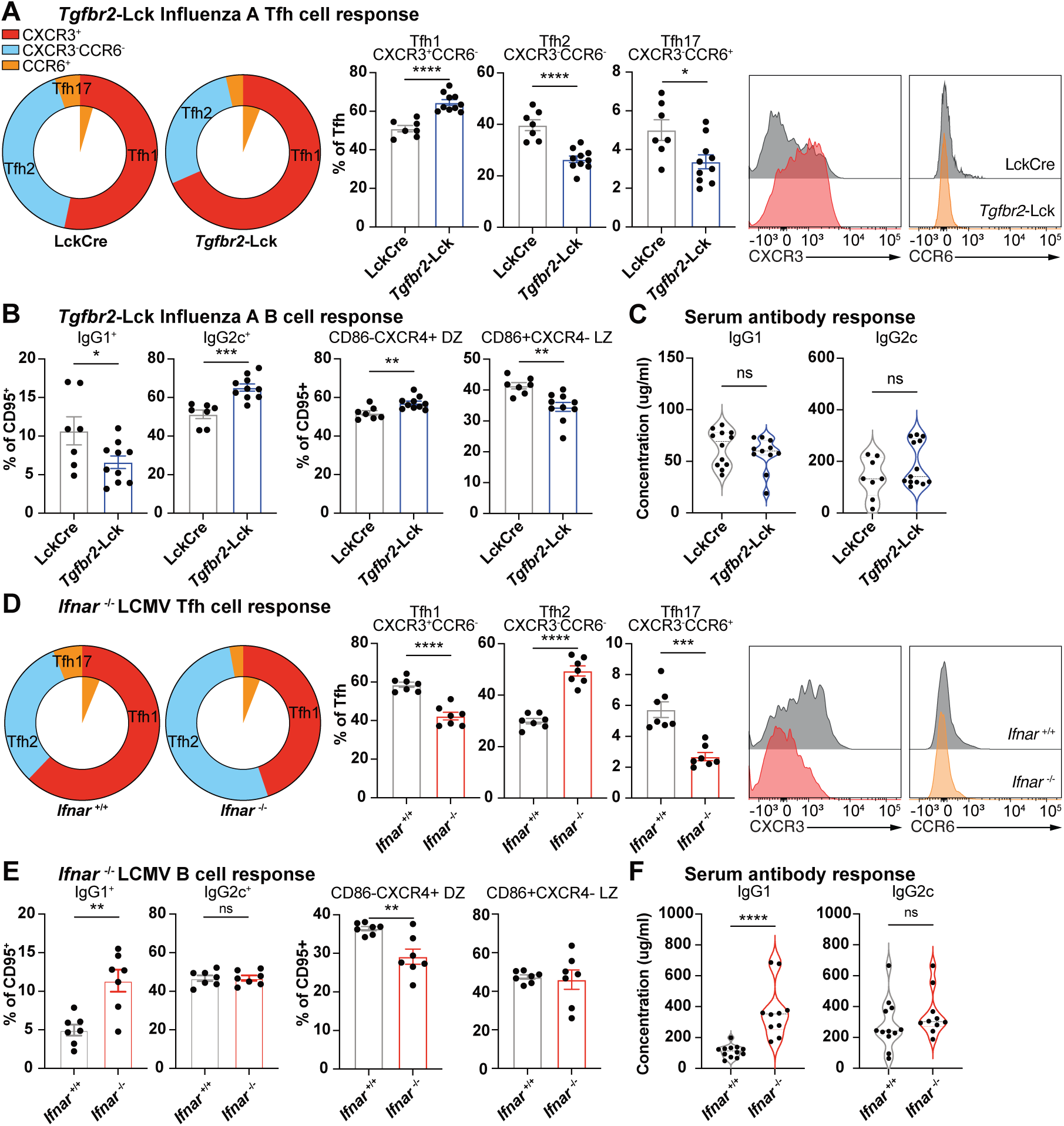
Pathogen-specific Tfh heterogeneity is directed by distinct cytokine signaling. (A to C) Draining lymph nodes analysis from *Tgfbr2*-Lck^Cre^ and Lck^Cre^ control mice infected with Influenza A virus. (A) CXCR3^+^ Tfh1, CXCR3^-^CCR6^-^ Tfh2, CCR6^+^ Tfh17 cells within the CD4^+^CD44^+^CXCR5^+^Ly6C^-^ Tfh population displayed as parts of whole, frequency of Tfh population, and representative histograms of CXCR3 and CCR6 expression in Tfh cells. Inner slice of pie chart displays CXCR3^+^CCR6^+^ co- expression. (B) IgG1 or IgG2c class switched B220^+^IgD^lo^CD95^+^ GC B cells and CD86^-^CXCR4^+^ dark zone (DZ) or CD86^+^CXCR4^-^ light zone (LZ) B cells from B220^+^IgD^lo^CD95^+^ GC B cells. (C) Serum IgG isotype concentration. (D to F) Draining lymph nodes analysis from *Ifnar* ^-/-^ and *Ifnar* ^+/+^ control mice infected with LCMV virus. (D) CXCR3^+^ Tfh1, CXCR3^-^CCR6^-^ Tfh2, CCR6^+^ Tfh17 cells within the CD4^+^CD44^+^CXCR5^+^Ly6C^-^ Tfh population displayed as parts of whole, frequency of Tfh population, and representative histograms of CXCR3 and CCR6 expression in Tfh cells. Inner slice of pie chart displays CXCR3^+^CCR6^+^ co-expression. (E) IgG1 or IgG2c class switched B220^+^IgD^lo^CD95^+^ GC B cells and CD86^-^CXCR4^+^ DZ or CD86^+^CXCR4^-^ LZ B cells from B220^+^IgD^lo^CD95^+^ GC B cells. (F) Serum IgG isotype concentration. Data show experiments of 7-10 mice per group and mean ± SEM. Statistical tests: Student’s *t* test.

### Evidence of pathogen-specific cytokine signaling in human tonsillar Tfh populations

Human tonsils are secondary lymphoid organs constantly exposed to the upper respiratory tract. Despite this, it has been proposed that minimal tonsillar Tfh heterogeneity exists, due to a lack of both transcriptional heterogeneity and identification of CXCR3^+^ and CCR6^+^ populations^11,54^. We therefore hypothesized that our derived pathogen-specific Tfh transcriptional signatures may provide new insight into Tfh heterogeneity within human lymphoid tissue. Paired scRNAseq and scCITEseq analysis was performed on Tfh cells (CD3^+^CD4^+^CD45RA^-^ CD45RO^+^CXCR5^+^) from tonsils of adults with sleep apnea but otherwise ostensibly healthy (Figure S8A). Previous studies have identified 2-5 transcriptionally distinct Tfh populations, based on their migration and differentiation states, rather than by functional capacity^91,92^. Here, unsupervised Louvain clustering of 24393 Tfh cells identified 11 transcriptionally distinct Tfh clusters (C) (Figure 6A). The ranked scores for pathogen-specific signatures (identified Figure 4) were overlayed onto UMAP pseudo-bulked clusters to indicate pathogen-driven Tfh subpopulations (Figure 6B). For all 3 individual tonsil samples, LCMV and Influenza signatures aligned to C8 and C11, respectively (Figures 6B to 6D). In addition to LCMV, *T. muris* also scored high onto C8, while *H. polygyrus* and *C. rodentium* signatures highlighted C3, C6 and C1, C2 respectively (Figures 6B to 6D). Dichotomy between clusters was observed with inverse alignment seen between C11 and C8 when ranked based on Influenza signature, between C3 and C11 when ranked based on *H. polygrus*, and between C1 and C3 when ranked based on *C. rodentium* (Figure 6B). As scRNAseq C8 and C11 separated furthest from other Tfh populations and were the most polarized by the pathogen-specific signatures, we performed further analysis using GSEA and visualized using vissE gene set enrichment analysis^93^. Consistent with the cytokine-guided differentiation of Tfh subpopulations, C8 was enriched in gene networks for IFN signaling, while C11 was enriched in gene networks for TGFβ signaling (Figure 6E). Similarly, GSEA-vissE analysis of C1 showed enrichment of cytokine signaling gene networks for IL-6 and IL-2 family cytokines, while C3 was enriched in IL-1 family signaling (Figure 6F). Thus, we reveal evidence that distinct cytokine pathways drive unique Tfh subpopulations in both mice and humans.

**Figure 6.**
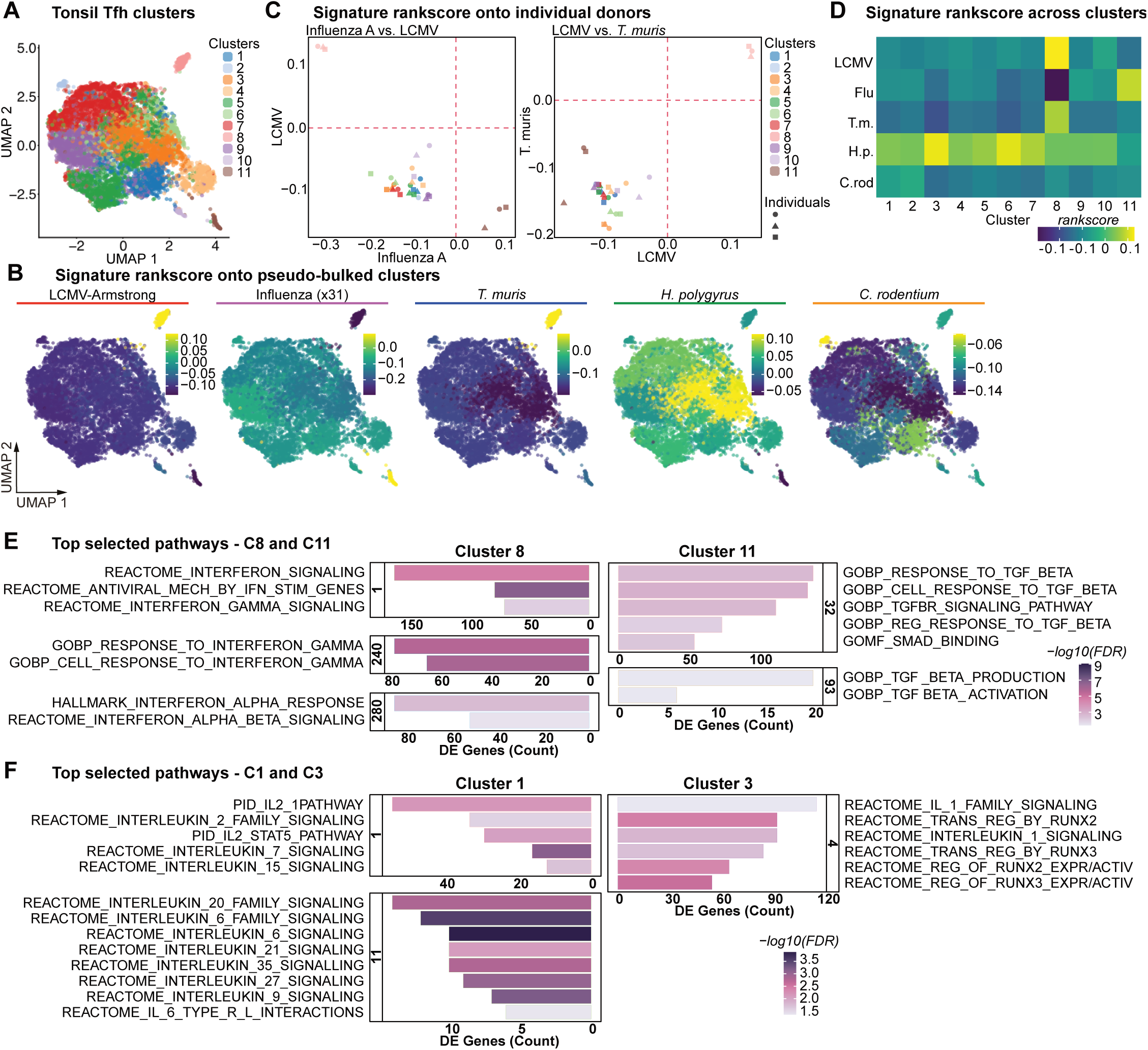
Evidence of diverse human tonsillar Tfh populations identified by unique cytokine signaling. scRNAseq of sorted CD3^+^CD4^+^CD45RA^-^CD45RO^+^CXCR5^+^CD27^+^ human tonsillar Tfh cells from three healthy adult donors. (A) UMAP dimensional reduction of data depicting 11 clusters based on louvain clustering using Jaccard similarity index (*k* = 10). (B) Overlay of mean ranked scores of each pathogen-specific signature onto UMAP clusters. Mean of the ‘*TotalScore’* from *singscore::simpleScore* function collated for each cluster. Pathogen-specific up- and down-regulated signature genes used. (C) Rank scores of pathogen-specific signatures for each pseudo-sample based on cluster and individual donors. (left) Influenza A vs LCMV (right) LCMV vs T.muris. (D) Heatmap of the mean rank scores of pathogen-specific signatures for each cluster. (E) Geneset Enrichment and vissE analyses identified individual clusters enriched in cluster 8 or 11 for the comparison Cluster 8 vs Cluster 11 and (F) individual clusters enriched in cluster 1 or 3 for the comparison Cluster 1 vs Cluster 3. Top gene sets of selected gene set clusters of interest shown as barplot of DE gene counts with corresponding gene statistics (FDR as color shade) from the DE analysis. GSEA was performed using Hallmarks c2 (’CP:REACTOME’, ’CP:PID’, ’CP:BIOCARTA’,’CP:KEGG’) and c5 (’GO:BP’,’GO:MF’) collections from the MsigDB, with false discovery rate (FDR) adjusted *p*≤0.05.

### Cell surface identification of subpopulations in human tonsil Tfh cells

As human cTfh subpopulations align with distinct B cell output^14,47–49^, we next sought to identify a tractable method to identify Tfh heterogeneity within human lymphoid tissue. For this we investigated the alignment of CITE-seq surface protein expression levels with the associated tonsil Tfh clusters. Key markers of cell identity showed concurrence between cell surface protein and mRNA expression for our Tfh cell gating strategy (CD4, CD45RO), and Tfh (CD279, CD278) and Teff (CD62L, KLRG1) core signatures (Figure 7A). Consistent with the lack of Tfh subpopulations defined by chemokine receptor expression, cell surface signal of CXCR3, CCR4, and CCR6 were minimally detected and did not indicate distinct Tfh clusters^11,54^ (Figure 7B). Instead, we identified several cell surface markers that discriminated either individual or groups of Tfh subpopulations (C1: CD43, CD99; C4: CD162; C8: CD71, CD151, CD134; C3:C7:C9: CD57, CD82 and C10: CD127) (Figure 7C). This approach paves the way to investigate how human Tfh subpopulations arise and directly tailor B cells in human health and disease.

**Figure 7.**
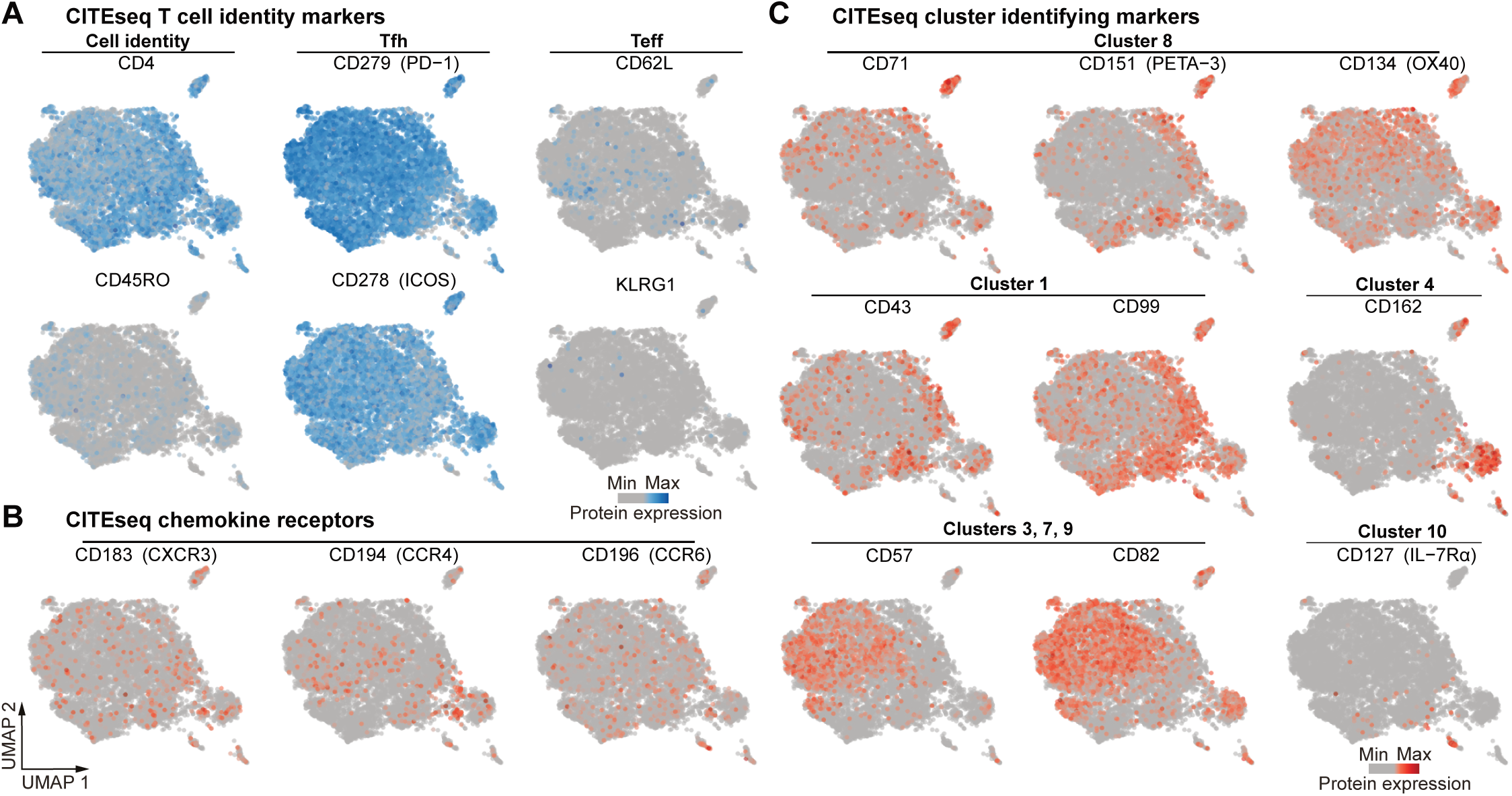
Cell surface proteins identify distinct human tonsillar Tfh subpopulations. scCITEseq surface protein expression of sorted CD3^+^CD4^+^CD45RA^-^ CD45RO^+^CXCR5^+^CD27^+^ human tonsillar Tfh cells from three healthy adult donors (as in Fig 7). scCITEseq surface protein expression (log counts) of (A) T cell identity markers, (B) chemokine receptors, and (C) cluster identifying markers, overlaid onto scRNAseq UMAP.

## DISCUSSION

The prevailing paradigm of Tfh differentiation has focused on bifurcation and balance between Tfh and individual Teff populations^15^. This model supports the lineage identity of Tfh cells^16,17,25^, yet fails to address how Tfh function is tailored depending on the inflammatory or infection setting. While transcriptional and functional Tfh heterogeneity has been observed, the breadth of heterogeneity or how it is established has not been formally tested. Here, we have taken a comparative approach, contrasting the transcriptional programing and function of Tfh cells that arise during diverse viral, helminth and bacterial infections. Our data is consistent with an initial bifurcation between Tfh and Teff cells (comprising of Th1, Th2 and Th17 populations). While the graded expression of T-bet correlated with Teff/Tfh ratios generated for each infection, the Tfh core signature was independent of *Tbx21* and the other lineage defining transcription factors, *Gata3* and *Rorc*. Instead, our approach indicated *Foxo1*, *Bach2*, and *Foxp1* likely oppose the Tfh program independently of inflammatory cues and pathogen class^67–73^. We confirm *Bcl6* expression as the transcriptional cornerstone of Tfh cells, regulating transcriptional regulators within both Tfh and Teff core programs^16–18,87,89,90^. Further, we identify several novel core transcriptional regulators that may additionally direct Tfh differentiation or function. Our data propose that a secondary differentiation branching occurs after or parallel with initial bifurcation, which directs the functional diversification of Tfh cells. At this step, Bcl-6 is co-expressed with distinct lineage defining transcription factors that superimpose a stimuli-specific program to direct function and tailoring of B cell responses (Figure S9). In addition to Teff and Tfh cells, equivalent transcriptional modules led by T-bet, GATA3 and RORγt exist for T regulator cells, innate lymphoid cells and dendritic cells^94–96^. Thus, our results highlight the dual transcriptional processes of lineage specification and functional transcriptional modules as central processes that underpin how the immune system adapts to diverse environmental challenges.

As Bcl-6 acts to repress alternative Teff fates, it remains unclear how opposing transcriptional regulators can be co-expressed within a single Tfh cell^16,17^. We propose that graded expression of lineage defining transcription factors navigate the tuning between lineage specification and functional tailoring. For example, it is striking that T-bet expression, despite varying with infection type, is always significantly higher in Teff compared to Tfh cells. Accordingly, interactions between Bcl-6 and T-bet alter recruitment to DNA loci to co-opt their transcriptional targets^29,43^. The concentration of cytokines, namely IL-2, IL-6 and IL-12, mediate this balance of Bcl-6, T-bet and GATA3 to direct differentiation to either Tfh or Teff fates^23,25,29–32,42,97^. Here we extend this concept to show local cytokine exposure specifies the functional quality of Tfh cells. We demonstrate that alterations of either TGFβ or IFN-I alter the differentiation of Tfh subpopulations. Notably, cytokine signaling appeared to be a stronger driver to shape Tfh heterogeneity than pathogen class, as Tfh in LCMV and *T. muris* featured IFN-I signaling, while Tfh cells in Influenza did not. These cytokine signaling pathways were confirmed in human Tfh populations, indicating their potential to drive the ontogeny of both Tfh heterogeneity in both human and mice. Accordingly, IFNAR deficiency mirrored both T-bet deletion in mice, and human T-bet loss-of-function mutations in humans^19,98^. There are likely other cytokines that additionally shape Tfh function. Indeed, *H. polygyrus* Tfh cells were not enriched for either IFN-I or TGFβ signaling, suggesting an alternate cytokine pathway tailors Tfh function in this context. While IL-4 is a candidate for this, this dataset is not available in the Hallmark MSigDB database. In human Tfh cells, we also identified clusters enriched for IL-6, IL-2 and IL-1 signaling. Thus, the dynamic local cytokine environment within draining lymph nodes, guides the generation of functionally distinct Tfh subpopulations. This model rationalizes multiple studies that have identified conflicting or context-dependent cytokines and transcription factors that modulate Tfh differentiation, suggesting that these may modify differentiation of specific Tfh subpopulations, rather than overall Tfh _fate_19,27,33,37,42,60,99.

Multiple questions regarding the ontogeny of Tfh heterogeneity remain to be investigated. To take an expansive view, here we analysed pathogen infections with a range of infection routes, lymph node drainage and varied antigen load and affinity. We propose that each of these likely plays a role in the outcome of Tfh function. Our dataset provides a resource for these aspects to be investigated individually.

Similarly, how the stability of individual Tfh subpopulations is maintained over the duration of the GC by distinct epigenetic regulation and how plasticity is regulated in memory and rechallenge remains to be determined. The combination of our mouse and human transcriptional dataset and human cell surface resources provide new tools for these questions to be addressed.

We demonstrate that cytokine-modified alteration of Tfh subpopulations in turn influenced GC B cell and antibody production. While our core signatures correlate with GC-Tfh populations, our data do not exclude that Tfh subpopulations influence antibody isotype usage prior to GC residency^100^. These observations offer new opportunities to target cytokine signaling pathways in order to direct B cell and antibody responses for the development of optimal humoral immunity in response to vaccines against a diverse range of pathogens. Further, identification of novel markers of human Tfh heterogeneity unlocks avenues to understand and treat antibody-mediated diseases such as immunodeficiency, allergy, asthma and autoimmunity.

## AUTHOR CONTRIBUTIONS

Conceptualization: J.R.G.

Methodology: L.D., C.W.T., A.A.S. and R.M.

Investigation: L.D., C.W.T., A.A.S., R.M., C.A., T.H., A.Z., L.C., A.K., L.H. and A.N.

Software and Formal Analysis: L.D. and C.W.T. Visualization: L.D. and C.W.T.

Resources: L.M., N.H., C.Z., N.L.LG., S.L.N., K.L.G-J., M.J.D., V.L.B. and J.R.G.

Writing – Original draft: L.D. and J.R.G. Funding Acquisition: J.R.G. Supervision: V.L.B. and J.R.G.

## ACKNOWLEDGMENTS

Cryopreserved tonsil samples were provided by Prof Cindy Ma (Garvan Institute). scRNAseq experiments were technically assisted by the WEHI Genomics Facility.

## STAR METHODS

**RESOURCE AVAILABILITY**

## Lead contact

Further information and requests for resources and reagents should be directed to and will be fulfilled by the lead contact, Joanna Groom (; @groomlab).

## Materials and Availability

This study did not generate new unique reagents.

## Data and code availability

Bulk RNA-seq data, single-cell RNA-seq data, and single-cell CITE-seq data have been deposited at GEO and are publicly available as of the date of publication. Accession numbers are listed in the key resources table.

All original code has been deposited at GEO and is publicly available as of the date of publication. DOIs are listed in the key resources table.

Any additional information required to reanalyze the data reported in this paper is available from the lead contact upon request.

## EXPERIMENTAL MODEL AND STUDY PARTICIPANT DETAILS

### Mice

Mice were bred and maintained on a C57BL/6 background under specific pathogen-free conditions. T-bet-ZsGreen reporter^61^, IL-4-AmCyan-IL-13-DsRed-IFN-γ-GFP reporter^8,63^, IL-21 GFP reporter^12^, FoxP3-RFP reporter^102^, *Tgfbr2*-Lck^Cre103^, and *Ifnar ^-/-^*^104^ mice have been previously described. All experiments were conducted in compliance with the Walter and Eliza Hall Institute Animal Ethics Committee and performed on sex-matched mice of 6-10 weeks of age.

### Infections

Mice inoculated intravenously with 3x10^3^ plaque-forming units (PFU) of LCMV-Armstrong were harvested 12 days post infection. Mice intranasally infected with 1x10^4^ PFU Influenza A virus strain HKx31 were harvested 10 days post infection. Mice infected via oral gavage with 2x10^9^ colony-forming units *Citrobacter rodentium*, 200 embryonated *Trichuris muris* eggs, or 200 L3 stage *Heligmosomoides polygyrus* larvae were harvested 12, 21 and 12 days post each respective infection.

### Human tonsil samples

Cryopreserved tonsil samples were obtained from three healthy adults (30-38 years-old; 1 female, 2 male). All procedures involving human participants were approved by and in accordance with the ethical standards of Human Research Ethics Committee at WEHI and the 1964 Helsinki Declaration and its later amendments.

## METHOD DETAILS

### Flow cytometry

All mice were harvested at the early peak of Tfh and GC B cell accumulation in the respective draining lymph nodes for each infection. Draining lymph node single-cell suspensions were stained for surface antigen expression using indicated antibodies for 20 minutes at 4°C, followed by viability dye staining for 10 minutes at 4°C. For cytokine detection, single-cell suspensions were stimulated in round-bottom tubes at 37°C + 5% CO2 in RPMI with 100 ng/ml of PMA (Sigma), 500 ng/ml Ionomycin (Sigma), 100 ng/mL Brefeldin A (BD Biosciences), 100 ng/mL monesin (BD Biosciences) and 10% FCS for 4 hours. Cytokine staining was performed using the BD Cytofix/Cytoperm Kit (BD Biosciences). Flow cytometry analysis was performed on a BD LSRFortessa X-20 and BD FACSymphony A3 Cell Analysers (BD Biosciences) and analyzed using FlowJo v10 (FlowJo LLC).

### Immunofluorescence staining and confocal microscopy

Lymph nodes were harvested and fixed in 4% Paraformaldehyde (Sigma) for 8 hours at 4°C, immersed in 30% sucrose overnight at 4°C, and embedded in OCT compound (Tissue-Tek). Tissues were cut via microtome (Leica) into 12-20μm sections and mounted onto Superfrost Plus slides. Slides were incubated in blocking buffer containing 2% Normal rat serum (Jackson ImmunoResearch) and 0.1% Triton X-100 (Sigma) in PBS for 24 hours at 4°C. Sections were stained with indicated antibodies in buffer containing 0.2% Normal rat serum (Jackson ImmunoResearch) and 0.01% Triton X-100 (Sigma) in PBS overnight at 4°C. Slides were washed by immersion in PBS containing 0.1% Triton X-100 (Sigma) and coverslips mounted with Prolong Diamond (ThermoFisher Scientific). Images were acquired on a LSM980 confocal microscope (Carl Zeiss MicroImaging) and processed using Zen Black (Zeiss) software.

### Serum Cytokine bead Array

Blood was collected from infected mice and centrifuged at 20,000*g* for 15 min. Serum supernatant was collected and loaded onto BD Cytometric Bead Array Mouse Th1/Th2/Th17 CBA kit (BD Biosciences), as per manufacturer’s instructions. Briefly, serum and mouse cytokine standards were added to cytokine capture beads and PE detection reagent in 96-well plate for 2 hours at room temperature, protected from light. The plate was washed two times in BD CBA wash buffer, resuspended and acquired on a BD LSRFortessa X-20 cell analyzer (BD Biosciences). Data analysis and standard curve generation was performed with FCAP Array Software version 3.0 (BD Biosciences).

### ELISA

High-binding 96-well ELISA plates (Sarstedt) were coated with relevant anti-IgG unconjugated antibody (Southern Biotech) and incubated overnight at 4°C followed by incubation with blocking buffer containing 1% bovine serum albumin (BSA) in PBS for 1 hour at room temperature. Supernatant was removed and plates were washed with PBS containing 0.04% Tween20 (ThermoFisher Scientific) followed by dH2O three times. Serum samples were diluted in 1% BSA in PBS, and serially diluted three-fold and incubated for 3 hours at 37°C. Plates washed three times, and incubated with secondary antibody conjugated to horseradish peroxidase (Southern Biotech) for 1.5 hours at 37°C. Plates were washed five times and OPD color reaction (Sigma) was added to each well. Absorbance was measured at 450nm using a FLUOstar Omega Microplate reader (BMG LABTECH). Average blank optical density (OD) values were subtracted from sample raw OD values.

### Preparation of draining lymph node samples for bulk RNA sequencing, data processing and analysis

#### Sample preparation and cell sorting

Single cell suspensions from draining lymph nodes were stained for surface antigen expression using indicated antibodies for 30 minutes at 4°C, followed by viability dye staining for 15 minutes at 4°C. Sample suspensions were sorted using BD FACSAria Fusion Flow Cytometer (BD Biosciences) to isolate Tfh cells.

#### Bulk RNA sequencing

RNA was isolated from sorted cells using RNeasy Plus Micro kit (QIAGEN) and library was prepared with SMART-seq PLUS kit (Takarabio). RNA sequenced using Illumina NextSeq 500 System on a HiSeq paired-end run. The R-subread aligner was used to align sequencing reads to the GRCm39 Mus musculus reference genome release version 103 and featureCounts^105^ was subsequently used to quantify transcripts per gene.

#### Data processing

Genes with low expression were filtered out using *edgeR*::*filterByExpr* function (v3.34.0)^106^, immunoglobin, TEC (To be experimentally confirmed) and gender related genes were removed. Samples with library size <1M reads were discarded. Relative log expression (RLE) plots and principal component analysis (PCA) were used to identify factors associated with the variations in the data and to identify batch effects. Outlier and samples with poor RNA quality were subsequently removed.

Normalization was then performed using the TMM method^107^. All bulk RNA-seq was performed in a single experiment, therefore no batch correction was performed.

#### Differential expression analysis

Differential expression analysis was performed using the *voom-limma* pipeline^108,109^ from the *limma* R/Bioconductor package (v3.48.0)^109^. Linear models were fitted using *limma::lmfit* to a design matrix accounting for biological factors of interest against the log expression of each transcript to identify differential expression between the contrast Tfh vs. Teff cells in either all infections or for individual infection. The t-tests relative to a threshold (TREAT) criterion was then applied^110^ to perform statistical tests and calculate the t-statistics, log-fold change (logFC). The statistical significance threshold was set to an adjusted *p*-value ≤0.05 and was calculated using the Benjamini Hochberg procedure.

#### Deriving the core Tfh signature

First, differential expression between the contrast Tfh vs. Teff cells from all infections was performed. Next, differential expression between the contrast Tfh vs. Teff cells for the five individual infections was performed. Genes that were intersecting directionally across 3 or more infections and present in the Tfh vs. Teff contrast from all infections were used for the core Tfh signature.

#### Deriving the pathogen-specific Tfh signatures

Pathogen-specific Tfh signatures were derived from differential expression results using the contrast Tfh cells from one infection vs. Tfh cells from all other infections.

#### Gene set enrichment analysis

The enrichment of marker gene sets from published Tfh and GC B cell datasets^64,82–84,88,111–113^ and precursor of exhausted and memory T cell^114–117^ datasets in the DE results was calculated using the fgsea package^118^ and visualized as normalized

enrichment score (NES) dot plots and as barcode plot of enrichment scores for the following gene sets^87–90,101^.

#### Visualisation of signatures

Differential expression between contrast Tfh vs. Teff cells from all infections visualized using ggplot2 R package (https://ggplot2.tidyverse.org). Visualization of the intersects from differential expression between contrast Tfh vs. Teff cells for the five individual infections performed using UpSetR package^119^. Signatures visualized on heatmaps using ComplexHeatmap R package^120^ for cell surface receptors and transcriptional regulators. Cell surface receptors genes were based on gene sets downloaded from Mouse Genome Database^121^ for cell surface (GO:0009986) and cell surface receptor signaling pathway (GO:0007166). Transcriptional regulators genes included transcription factors annotated in Immunological Genome Project (ImmGen) database ^122^, mouse tissue transcription factor atlas^123^, and gene sets downloaded from Mouse Genome Database for transcription factor activity (GO:0003700), transcription activator activity (GO:0001216), transcription repressor activity (GO:0001217), and transcription factor binding (GO:0008134). Bcl-6 network transcriptional regulator genes in Tfh core (without cell cycle genes) were visualized for Tfh vs. Teff cells and Tfr vs. Teff cells contrasts using ggplot2 R package.

### Preparation of human tonsil samples for cell sorting, single cell RNA sequencing, and CITE sequencing

#### Sample preparation and cell sorting

Cryopreserved mononuclear cell suspensions from tonsil samples were thawed and processed to form single cell suspensions. Samples were stained for surface antigen expression using indicated antibodies and TotalSeq HashTags (BioLegend) for 30 minutes at 4°C, followed by viability dye staining for 15 minutes at 4°C. Sample suspensions were sorted using BD FACSAria Fusion Flow Cytometer (BD Biosciences) to isolate Tfh cells.

#### Single cell RNA sequencing and CITE sequencing

Equal numbers of sorted cells from each tonsil sample were pooled and stained with TotalSeq-A Human Universal Cocktail in accordance with manufacturer’s protocols. Sequence tags of DNA-barcoded mAbs for CITE-seq were incorporated into existing scRNA-seq to enable simultaneous quantification of protein and gene expression by single cells. 40,000 cells from the stained Tfh pool underwent 3’-tag-capture using the 10X Genomics Chromium Controller. Library preparation was conducted in preparation for sequencing according to manufacturer’s protocols. Next Generation Sequencing was performed using an Illumina NextSeq 500 System and Illumina NovaSeq 6000 System.

#### Data processing and demultiplexing of single cell RNA sequencing data

For data processing, reads from each capture were processed using 10X Genomics Cell Ranger software (v7.0.0). Firstly, ‘cellranger mkfastq’ and bcl2fastq (v2.19.1) were used to convert and demultiplex the Illumina sequencer’s BCL files into FASTQ files for each of the gene expression (GEX), antibody derived tag (ADT), and hashtag oligo (HTO) libraries. Secondly, ‘cellranger multi’ was used with default settings to generate count matrices. The GEX data were mapped and quantified against the 10X Genomics pre-built mm10 (GENCODE vM23/Ensembl 98) reference groups and transcriptome (2020-A (July 7, 2020) version) and the feature barcoding (ADT and HTO) data were quantified against a ‘feature.csv’ file containing the barcode sequences provided by BioLegend. Finally, the DropletUtils R/Bioconductor package (v1.18.1) was then used to load the Cell Ranger output files into R (v4.2.1) and to identify nonempty droplets using ‘emptyDrops’ method with default settings^124^. For demultiplexing, the ‘demuxmix’ method from the demuxmix (v1.0.0) R/Bioconductor package^125^ was applied to the HTO data, with the ‘naive’ model and default parameters, to multiplex non-empty droplets to their sample of origin. This was performed separately for each capture. A droplet was assigned to a sample if the best demuxmix assignment matched the corresponding HTO combination of a sample or was otherwise assigned as a ‘multiplet,’ ‘negative,’ or ‘uncertain’ sample.All scripts used will be made available upon manuscript acceptance from https://github.com/WEHISCORE/G00 _Dalit.

### Analysis of single cell RNA sequencing data

#### Quality Control, clustering and phenotype scoring

The demultiplexed scRNA-seq data was assessed where low quality cells, cells with high mitochondrial genes (≥ 20%), doublets and cells with unknown HTO tags were removed, leaving 23126 cells. The data was then normalized using R packages *scran*^126^ and *scuttle*^127^, converted to log-normalised counts and the top 2000 Highly variable genes (HVGs) were derived and used to run principal component analysis (PCA) dimension reduction. Data was clustered using the louvain method based on a shared nearest neighbour graph for k=10 nearest neighbors and the jaccard weighting scheme (using *igraph* and *scran*), resulting in 11 clusters.

The Tfh phenotypic states of the cells were investigated by computing the Tfh and pathogen specific Tfh signatures (based on the signatures derived from the bulk RNA-seq analysis) using the R package singscore^128,129^. Scores for the Tfh signatures are visualised by overlaying across the UMAP using *scater::plotReducedDim* function.

#### Differential expression analysis

Differential expression (DE) was conducted using a pseudo-bulked approach based on “clusters” and “samples” using aggregateAcrossCells function in scuttle^127^, and after filtering out samples with less than 10 cells and applying gene level QC using *edgeR::filterByExpr*, leaving 31 pseudo-samples and 12238 genes for analysis.

Differential expression analyses were conducted using a *voom-limma- duplicatecorrelation* pipeline via the *edgeR::voomLmFit* function to fit a linear model with “cluster” as the covariate, and to estimate the consensus correlation across mice and account for mice variation as a random effect^106^. DE expression was conducted for the following comparisons based on clusters identified from the pathogen-specific signature rankscore analysis: A) Cluster 8 vs Cluster 11 and B) Cluster 1 vs Cluster 3. An empirical Bayes moderated t-statistic was generated with multiple testing adjustment carried out using the Benjamini–Hochberg procedure to identify statistically significant genes (adjusted P < 0.05).

#### VissE gene set network analysis

Gene set enrichment analyses (GSEA) were performed based on the DE results using the fry approach from the *limma* package based on the gene sets downloaded from Molecular Signature Database ^130^ categories Hallmarks, Curated (c2: Reactome, PID, Biocarta and KEGG) and Ontology (c5: BP and MF). Gene sets with a false discovery rate (FDR) less than 0.05 were considered significant. Significant genesets were analyzed using vissE ^93^ R package, clustering similar processes and identifying overarching biological themes. The gene set network was generated using vissE by applying an Overlap Coefficient threshold of 0.25 and 0.15 (Up- and Down-regulated respectively) for both the Cluster 8 vs Cluster 11 and Cluster 1 vs Cluster 3 comparisons respectively. Clusters were identified using the walktrap algorithm and clusters with less than 2 gene sets were removed. Gene sets clusters that were enriched for each comparison were identified alongside the gene statistics and common biological themes highlighted.

#### Data processing and visualisation of CITE-seq data

CITE-seq data was extracted and normalized using the CLR (centered log ratio transformation) method across cells from the *Seurat::NormalizeData* function ^131^. The logcount expression for selected markers were visualized by overlaying onto the scRNA-seq UMAPs of tonsil Tfh.

### Antibodies and dyes for staining

#### T cell analysis

Single cell suspensions were stained with fixable viability stain 700 (1:1,000; BD Biosciences); anti-CD4 (1:600; clone GK1.5; BD Biosciences); anti-CD3 (1:200; clone 145-2C11; BD Biosciences); anti-CD44 (1:200; clone IM7; BD Biosciences); anti-Ly6C (1:400; clone HK1.4; BioLegend); anti-CXCR5 (1:200; clone L138D7; BioLegend); anti-CD162 (1:800; clone 2PH1; BD Biosciences); anti-PD-1 (1:200; clone RMP1-30; BioLegend); anti-CD62L (1:400, clone MEL-14; Thermo Fisher); anti-CD127 (1:200; clone SB/199; BD Biosciences); anti-IFNγ (1:400; clone XMG1.2; BD Biosciences); anti-IL-4 (1:400; clone 11B11; BD Biosciences); anti-IL-17A (1:400; clone TC11-18H10; BD Biosciences); anti-CXCR3 (1:200; clone CXCR3-173; BioLegend) and anti-CCR6 (1:200; clone 29-2L17; BioLegend).

#### B cell analysis

Single cell suspensions were stained with fixable viability stain 700 (1:1,000; BD Biosciences); anti-B220 (1:800; clone RA3-6B2; BD Biosciences); anti-CD138 (1:400; clone 281-2; BD Biosciences); anti-CD95 (1:600; clone JO2; BD Biosciences); anti-IgD (1:200; clone 11-26c; WEHI Antibody Facility); anti-CD38 (1:600; clone 90; eBioscience); anti-CD86 (1:200; clone GL1; BD Biosciences); anti- CXCR4 (1:200; clone 2B11; Invitrogen); anti-IgG1(1:200; clone X56; BD Biosciences) and anti-IgG2a/2b (1:200; clone R2-40; BD Biosciences).

#### Single cell sorting for mouse bulk RNAseq

Single cell suspensions were stained with fixable viability stain (1:1,000; BD Biosciences); anti-CD4 (1:600; clone GK1.5; BD Biosciences); anti-CD44 (1:200; clone IM7; BD Biosciences); anti-CXCR5 (1:200; clone L138D7; BioLegend) and anti-PD-1 (1:200; clone RMP1-30; BioLegend).

#### Single cell sorting of human tonsil

Single cell suspensions were stained with fixable viability stain (1:1,000; BD Biosciences); anti-CD3 (1:20; clone SK7; BD Biosciences); anti-CD4 (1:20; clone SK3; BioLegend); anti-CD8 (1:20; clone SK1; BioLegend); anti-CD45RA (1:50; clone HI100; Invitrogen) and anti-CD45RO (1:20; clone UCHL1; BioLegend).

#### Confocal

Lymph node tissue sections were stained with anti-CD4 (1:100; clone GK1.5-7; WEHI Antibody Facility); anti-IgD (1:200; clone 11-26c; eBioscience) and anti-GL7 (1:100; clone GL7; Biolegend).

## QUANTIFICATION AND STATISTICAL ANALYSIS

### Quantification and statistical analysis

Statistical differences between groups in datasets with one categorical variable were evaluated by unpaired t-tests (2 groups) or one-way ANOVA (more than 2 groups) corrected for multiple comparisons. Statistical differences between groups in datasets with two categorical variables were evaluated by two-way ANOVA corrected for multiple comparisons. P < 0.05 was considered statistically significant.

* P < 0.05; ** P < 0.01; *** P < 0.001; *** P < 0.0001. All experimental data are presented as mean ± standard error of the mean (s.e.m.) with statistical analysis performed using Prism 9 (GraphPad Software).

---

**Figure S1.**
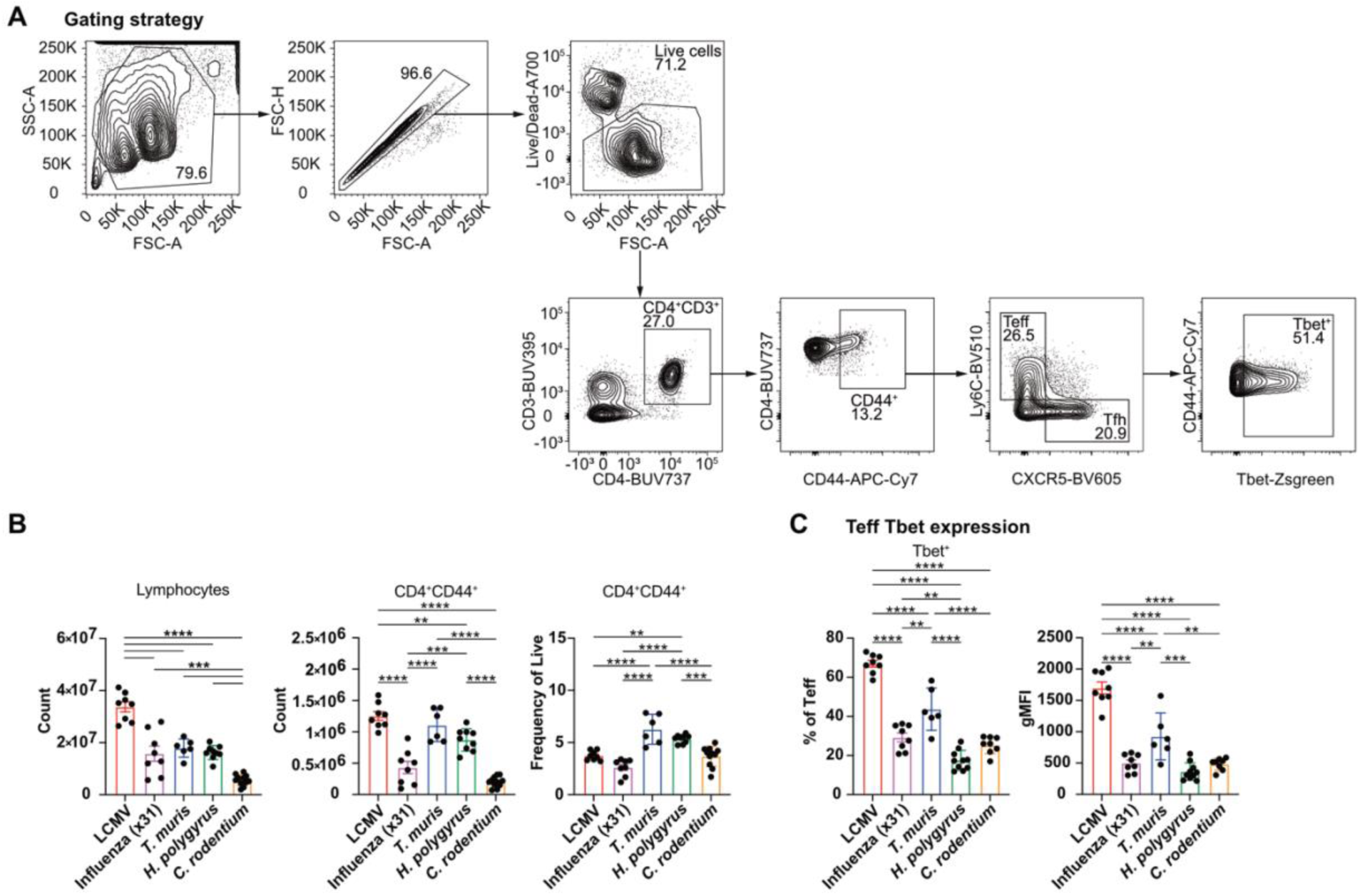
Diverse infections induce Teff populations with high T-bet expression. (A) Gating strategy to identify Tfh and Teff cell populations and ZsGreen_T-bet reporter expression. (B to C) Analysis of draining lymph node cells from wildtype and ZsGreen_T-bet reporter mice infected with indicated pathogens at early peak GC response. (B) Total lymphocyte counts, CD4^+^CD44^+^ cell counts and frequency of CD4^+^CD44^+^ cells of live cells. (C) ZsGreen_T-bet reporter^+^ frequency and gMFI of CD4^+^CD44^+^CXCR5^-^Ly6C^+^ Teff cells. Data show experiments of 6-10 mice per group and mean ± SEM. Statistical tests: one-way ANOVA of multiple comparisons.

**Figure S2.**
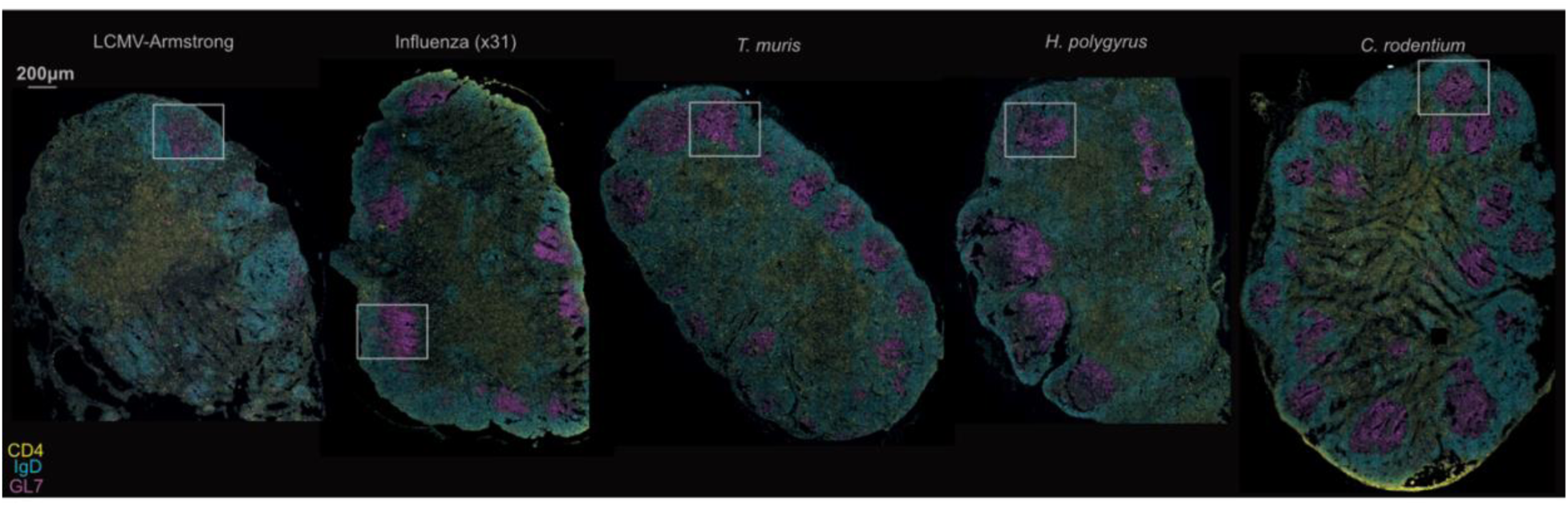
Conserved GC morphology in draining lymph nodes following infection with diverse pathogens. Immunofluorescent staining of draining lymph node of infected ZsGreen_T-bet reporter mice. Boxes indicate zoom of GC shown in Fig. 1F. Data are representative of 2-3 mice per group. Yellow: CD4, blue: IgD, magenta. Scale bar consistent for all images, 200μm.

**Figure S3.**
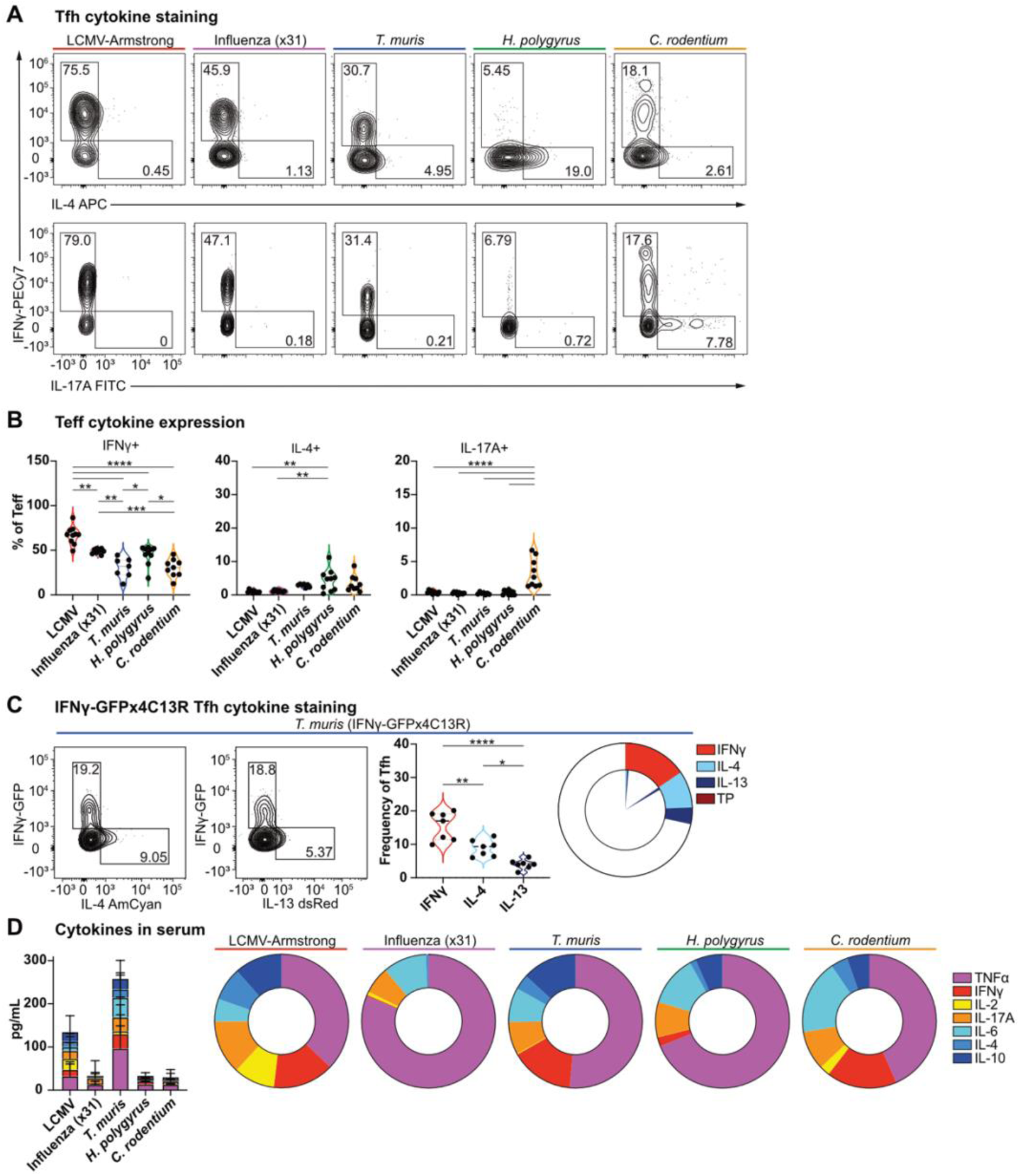
Heterogenous Tfh populations display diverse cytokine profiles reflected, similar to that of Teff and systemic cytokine milieu. Analysis of (A to C) draining lymph node cells and (D) serum from (A, B and D) wildtype and (C) IFNγ-GFPx4C13R reporter mice infected with indicated pathogens at early peak GC response. (A) Representative plots of Tfh cell IFNγ, IL-4, and IL-17 cytokine production. (B) Frequency of CD4^+^CD44^+^CXCR5^+^Ly6C^-^ Teff cell produced IFNγ, IL-4, IL-17A. (C) IFNγ-GFPx4C13R reporter expression in CD4^+^CD44^+^CXCR5^+^Ly6C^-^ Tfh cells following *T. muris* infection. Representative plots, frequency of cytokine reporter with Tfh population and displayed as parts of whole Tfh population. Inner slice displays cytokine co-expression. (D) Serum cytokine concentration (pg/mL) in serum. Cytokines displayed as parts of total cytokines analyzed. Data show experiments of 7-10 mice per group and mean ± SEM. Statistical tests: one-way ANOVA of multiple comparisons.

**Figure S4.**
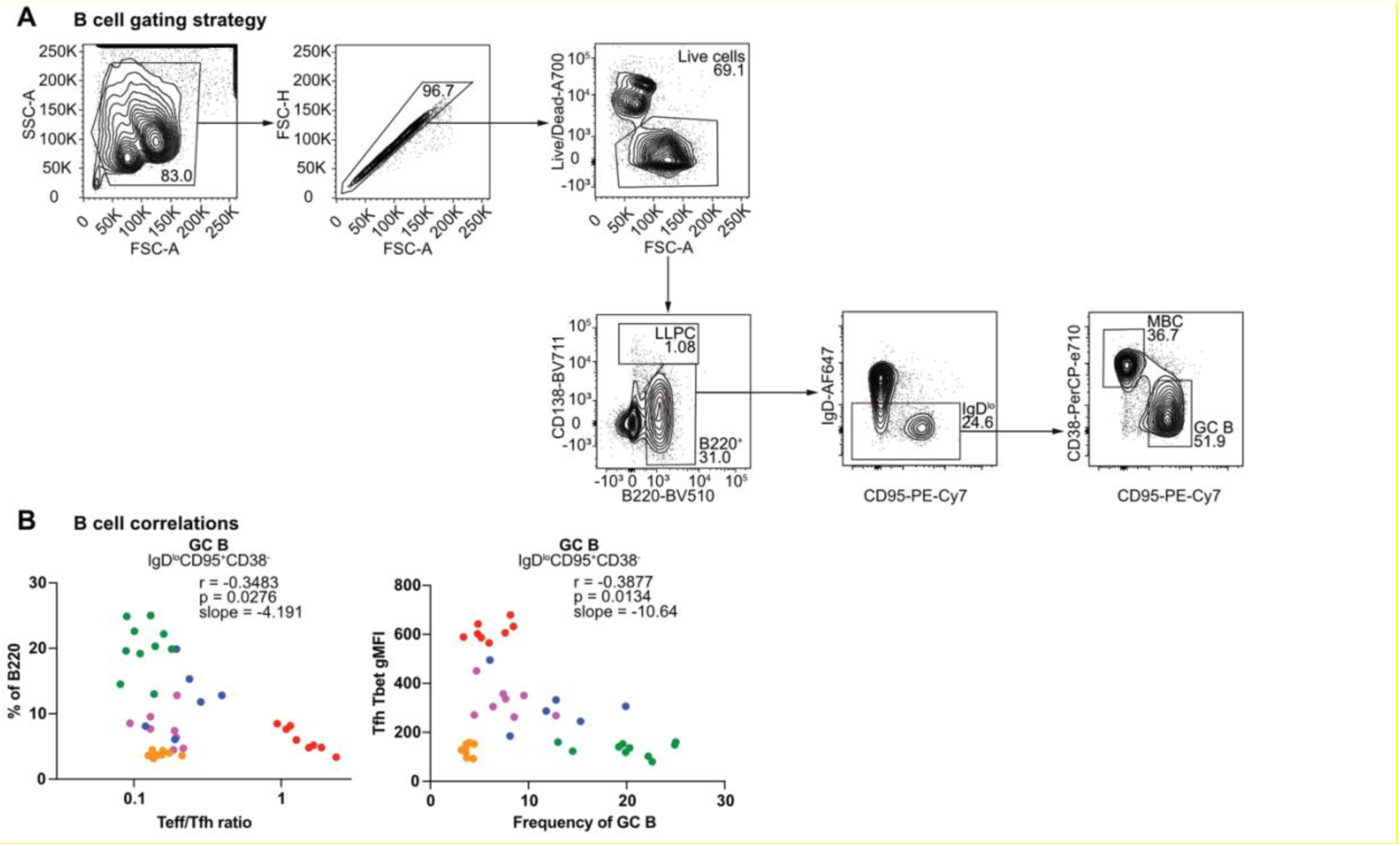
Pathogen-induced Tfh heterogeneity correlates tailored B cell responses. (A) Gating strategy to identify GC and MBC populations. (B) Analysis of B220^+^IgD^lo^CD95^+^CD38^-^ GC B cells from ZsGreen_Tbet reporter mice infected with indicated pathogens at early peak GC response. (B) Correlation of frequency of GC B to the ratio of Teff/Tfh cells and to ZsGreen_T-bet reporter expression gMFI across infections. Data show experiments of 6-10 mice per group. Statistical tests: Pearson correlation of two-tailed P value.

**Figure S5.**
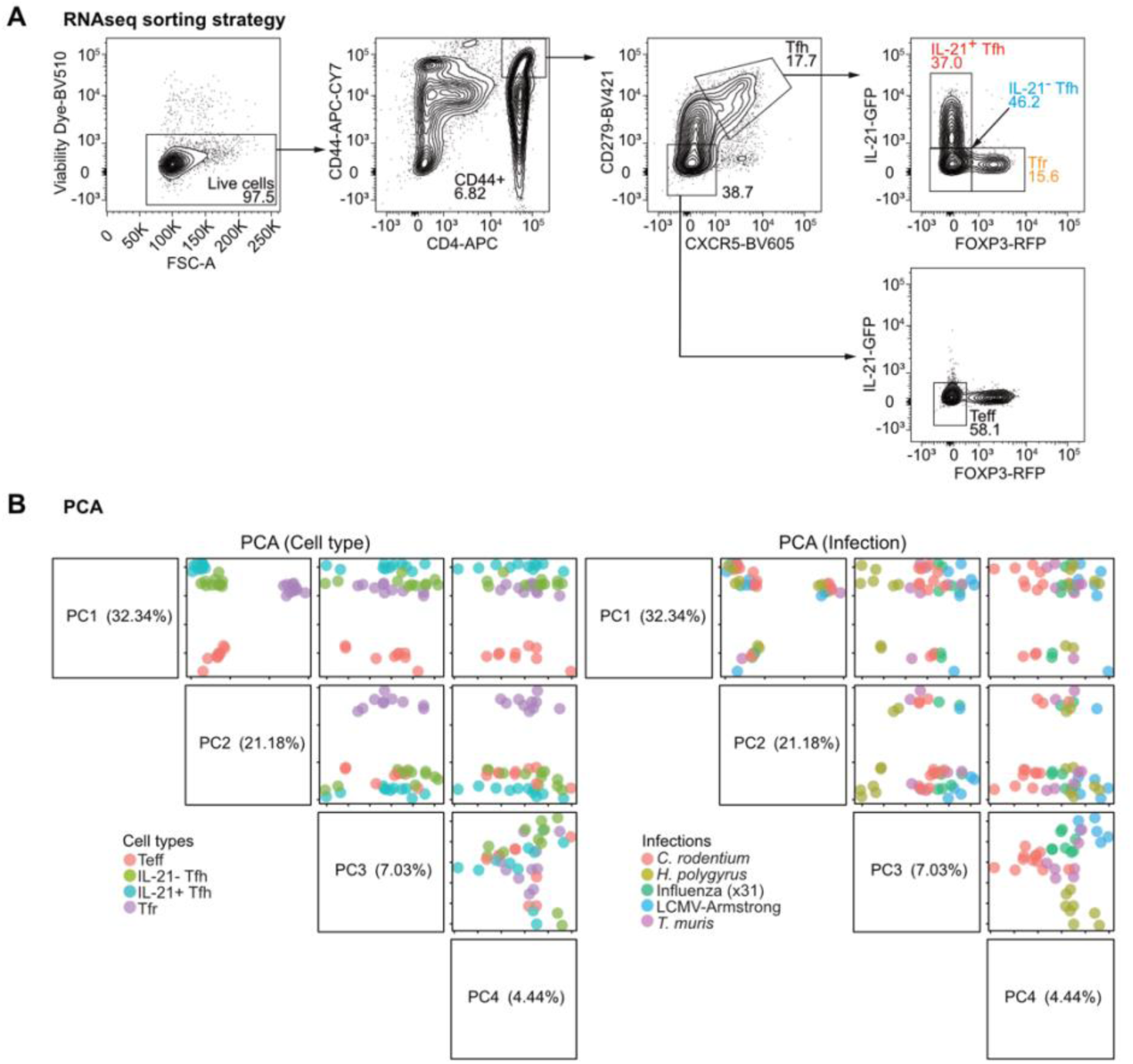
Pathogen-specific transcriptional programs separate by cell type and infection. (A to B) RNAseq of CD4^+^CD44^+^PD1^+^CXCR5^+^FoxP3-RFP^-^IL-21-GFP^+^ Tfh, CD4^+^CD44^+^PD1^+^CXCR5^+^FoxP3-RFP^-^IL-21-GFP^-^ Tfh, CD4^+^CD44^+^PD1^+^CXCR5^+^FoxP3- RFP^+^IL-21-GFP^+^ Tfr, and CD4^+^CD44^+^PD1^-^CXCR5^-^FoxP3-RFP^-^IL-21-GFP^-^ Teff cells from draining lymph nodes of wildtype mice infected with indicated pathogens at early peak GC response. (A) Gating strategy of sorted populations. (B) Principal Component Analysis (PCA) plots of sorted IL-21^+^ Tfh, IL-21^-^ Tfh, Tfr, and Teff cell transcriptomes stratified by either (left) cell type or (right) infection type. Data show independent samples of 2-3 per cell type per infection.

**Figure S6.**
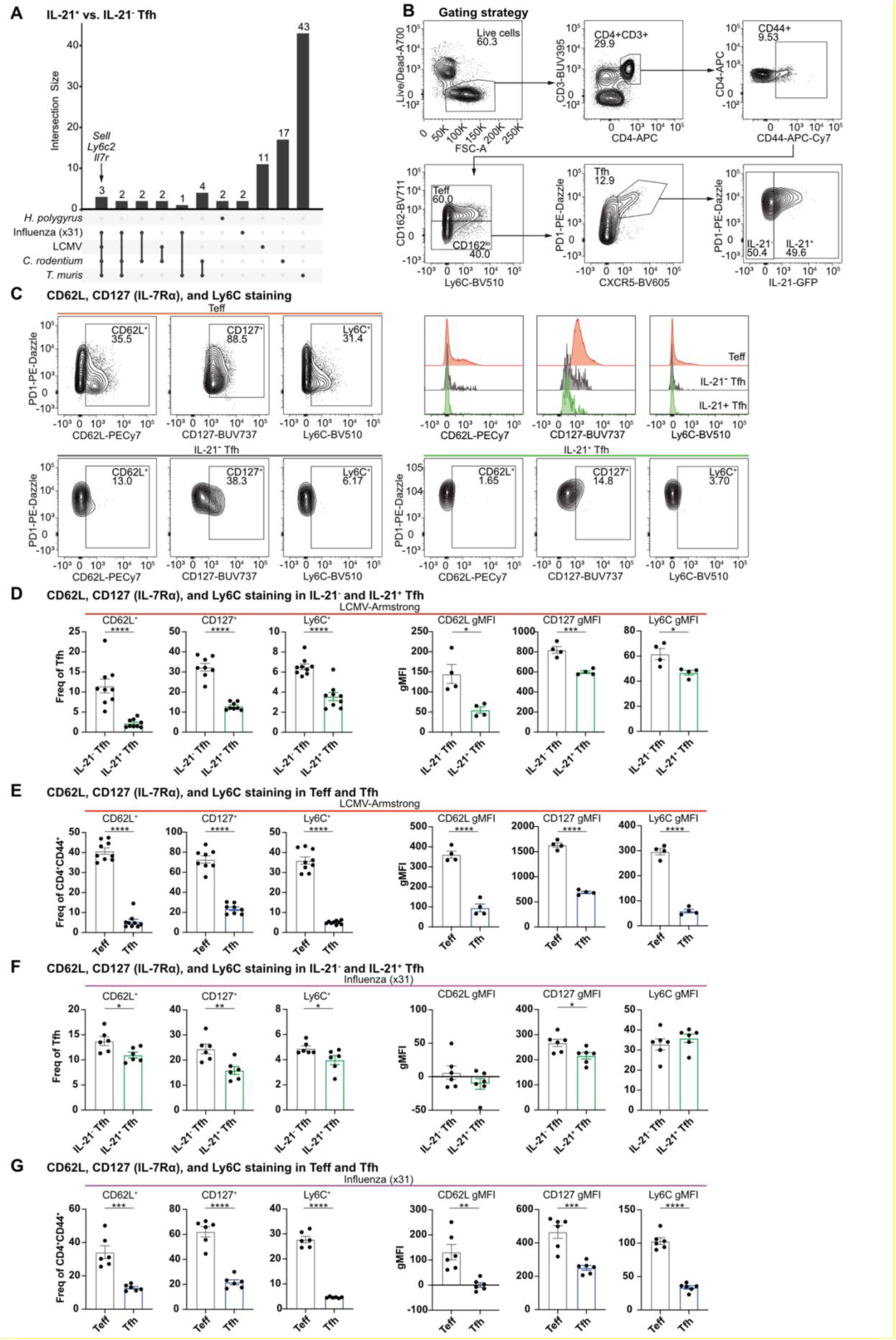
IL-21^+^ Tfh and IL-21^-^ Tfh cells are transcriptionally similar across diverse infections. (A) Upset plot displaying intersects of genes expressed differentially between IL-21^+^ Tfh and IL-21^-^ Tfh cells from separate infections in RNAseq analysis of CD4^+^CD44^+^PD1^+^CXCR5^+^FoxP3-RFP^-^IL-21-GFP^+^ Tfh and CD4^+^CD44^+^PD1^+^CXCR5^+^FoxP3-RFP^-^IL-21-GFP^-^ Tfh cells as in Fig 2. Three genes intersecting across 4 infections indicated with arrow. (B to G) Analysis of CD4^+^CD44^+^CXCR5^+^CD162^lo^ Tfh cells and CD4^+^CD44^+^ CXCR5^-^CD162^hi^ Teff cells from draining lymph nodes from wildtype mice infected with indicated pathogens, LCMV-Armstrong or Influenza A at early peak GC response. (B) Gating strategy to identify Teff, IL-21^-^ Tfh, and IL-21^+^ Tfh cells. (C) Representative plots and histograms of CD62L, CD127, and Ly6C staining on IL-21^-^ Tfh, IL-21^+^ Tfh cells and Teff cells in LCMV infection. (D to G) Frequency and gMFI of CD62L^+^, CD127^+^, and Ly6C^+^ on (D and F) IL-21^-^ Tfh, IL-21^+^ Tfh cells, frequency of total Tfh population (E and G) Tfh and Teff, frequency of CD4^+^CD44^+^ population. (D to E) in LCMV infection, (F to G) in Influenza A infection. Data show experiments of 4-6 mice per group and mean ± SEM. Statistical tests: Student’s *t* test.

**Figure S7.**
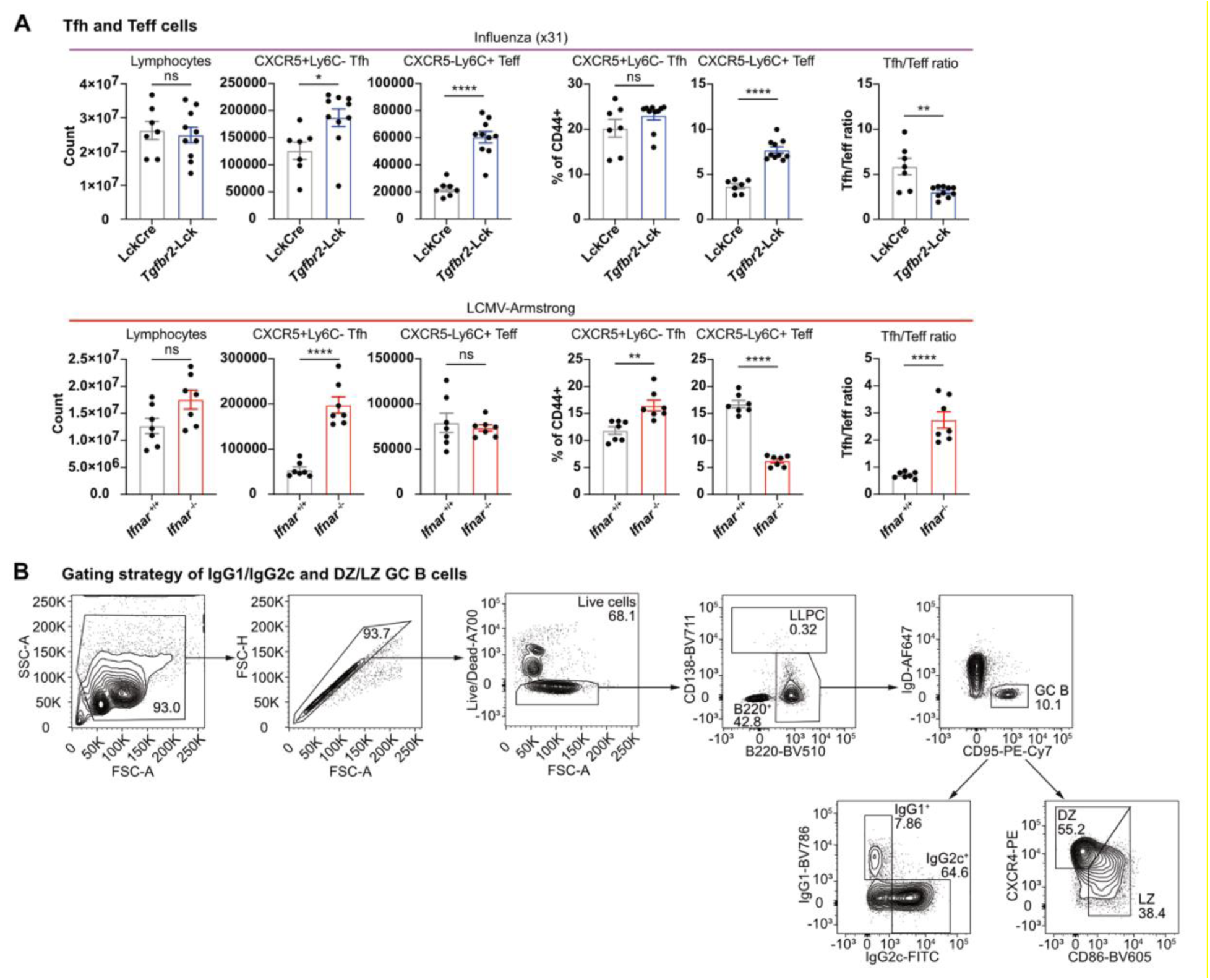
Pathogen-induced cytokine signaling influence Tfh and Teff bifurcation, Tfh subpopulations, and Tfh cells capacity to direct B cell and antibody output. (A) Draining lymph nodes analysed from either *Tgfbr2*-Lck^Cre^ and Lck^Cre^ control mice infected with Influenza A virus, or *Ifnar* ^-/-^ and *Ifnar* ^+/+^ control mice infected with LCMV at early peak GC response. (A) Total lymphocyte counts, CD4^+^CD44^+^CXCR5^+^Ly6C^-^ Tfh and CD4^+^CD44^+^CXCR5^-^Ly6C^+^ Teff cell counts, and Tfh and Teff cell frequencies. (B) Gating strategy to identify IgG1 and IgG2 class switched GC B cells, and dark zone (DZ) and light zone (LZ) GC B cells. Data show experiments of 7-13 mice per group and mean ± SEM. Statistical tests: Student’s *t* test.

**Figure S8.**
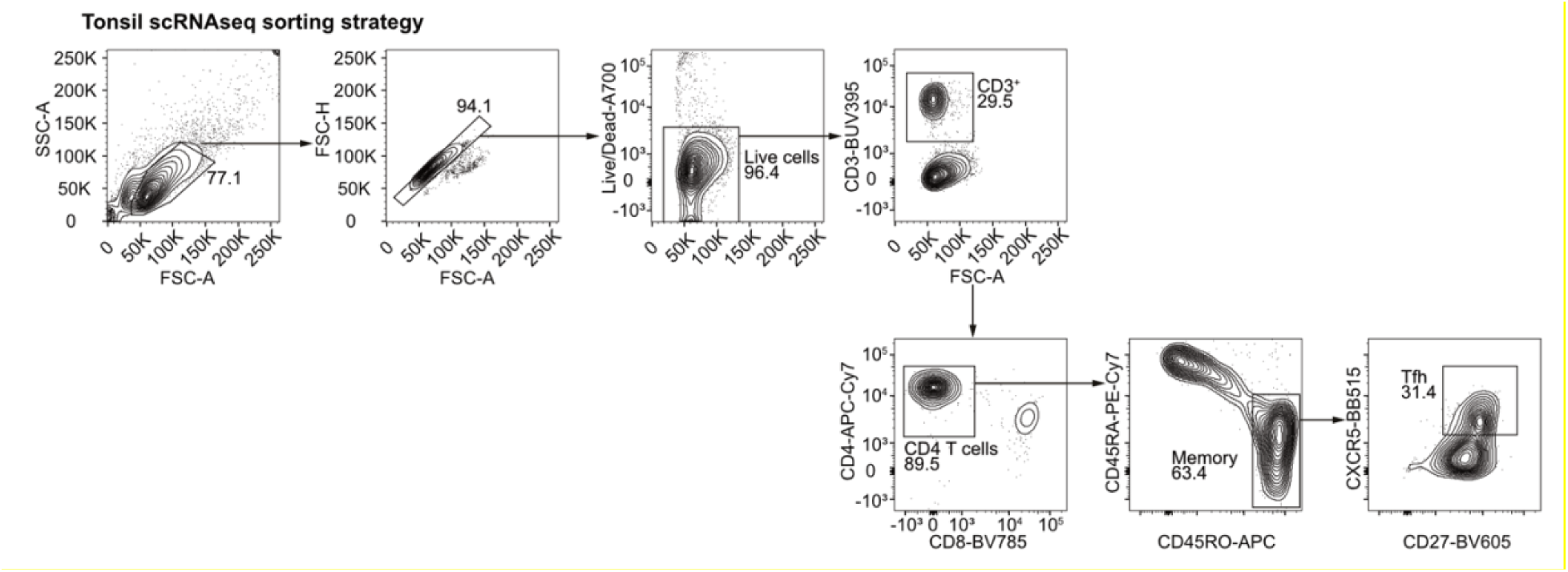
Human tonsillar Tfh population sorting gates. Gating strategy to sort CD3^+^CD4^+^CD45RA^-^CD45RO^+^CXCR5^+^CD27^+^ human tonsillar Tfh cells from three healthy adult donors for paired scRNA-seq and CITE-seq.

**Figure S9.**
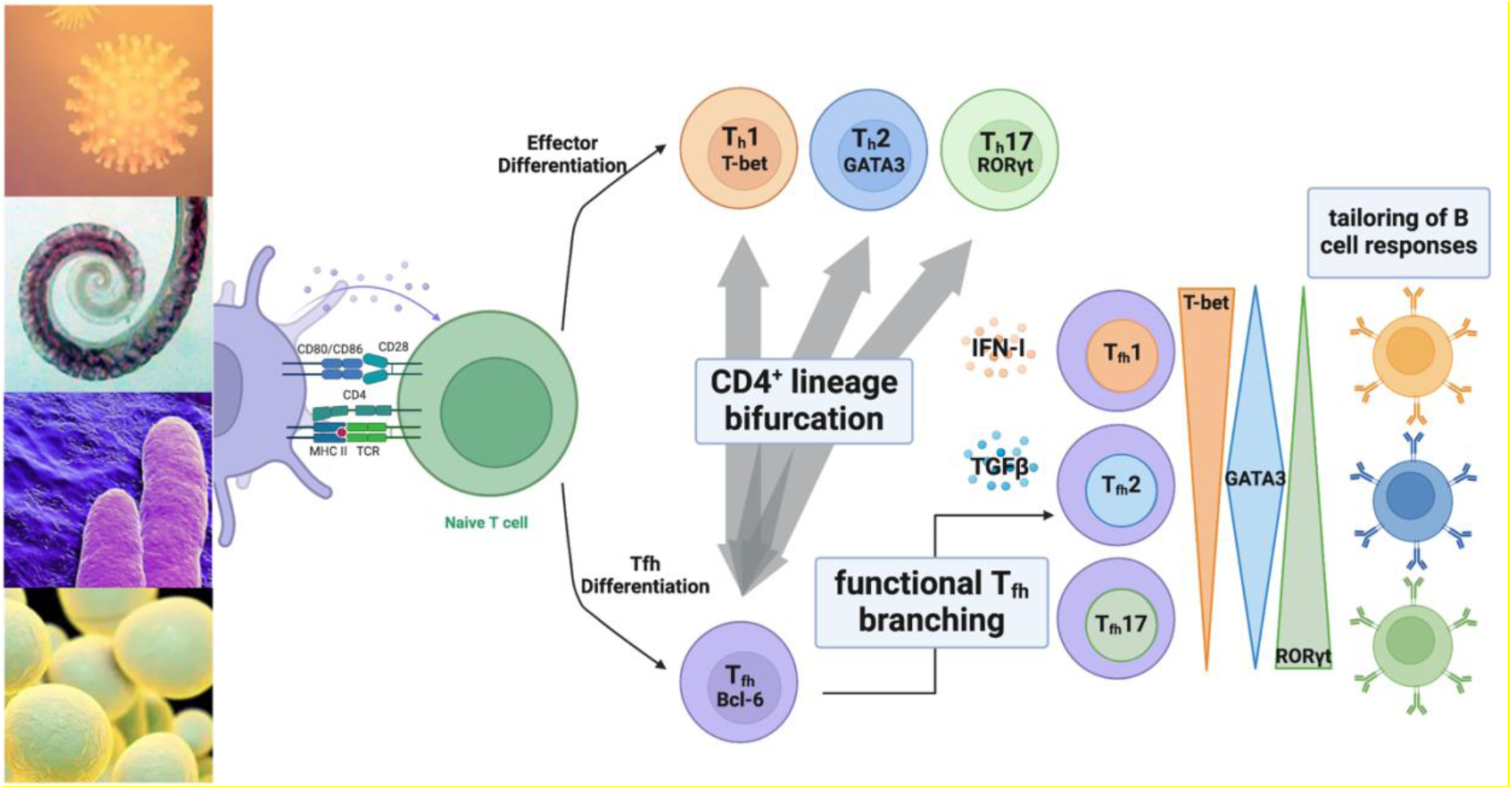
Extension of bifurcation model for Tfh differentiation to account for stimuli-specific functional heterogeneity. Environmental cues from viral, helminth, bacterial and fungal pathogens are integrated to drive heterogeneous CD4^+^ T cell differentiation and functional capacity. Tfh lineage specification is determined by Bcl-6 in contrast to Teff populations (Th1, Th2, Th17). Local integration of cytokines, including IFN-I and TGFβ leads to graded expression of the lineage defining transcription factors, T-bet, GATA3 and RORγt within the Tfh lineage. Tfh subpopulations in turn influence the quality of B cell responses and antibody production.

